# Investigating the effects of chimerism on the inference of selection: quantifying genomic targets of purifying, positive, and balancing selection in common marmosets (*Callithrix jacchus*)

**DOI:** 10.1101/2025.07.01.662532

**Authors:** Vivak Soni, Cyril J. Versoza, Susanne P. Pfeifer, Jeffrey D. Jensen

**Author notes:** co-corresponding authors; jointly supervised the project (;).

## Abstract

The common marmoset (*Callithrix jacchus*) is of considerable biomedical importance, yet there remains a need to characterize the evolutionary forces shaping empirically observed patterns of genomic variation in the species. However, two uncommon biological traits potentially prevent the use of standard population genetic approaches in this primate: a high frequency of twin-births and the prevalence of hematopoietic chimerism. Here we characterize the impact of these biological features on the inference of natural selection, and directly model twinning and chimerism when performing inference of the distribution of fitness effects to characterize general selective dynamics as well as when scanning the genome for loci shaped by the action of episodic positive and balancing selection. Results suggest a generally increased degree of purifying selection relative to human populations, consistent with the larger estimated effective population size of common marmosets. Furthermore, genomic scans based on an appropriate evolutionary baseline model reveal a small number of genes related to immunity, sensory perception, and reproduction to be strong sweep candidates. Notably, two genes in the major histocompatibility complex were found to have strong evidence of being maintained by balancing selection, in agreement with observations in other primate species. Taken together, this work, presenting the first whole-genome characterization of selective dynamics in the common marmoset, thus provides important insights into the landscape of both persistent and episodic selective forces in this species.

## INTRODUCTION

The common marmoset (*Callithrix jacchus*) is a platyrrhine native to east-central Brazil (Rylands and Faria 1993; Rylands et al. 2009; Garber et al. 2019), and a prominent model in biomedical research, with a growing emphasis on aging, gene editing / therapy, neuroscience, and stem cell research in recent years (Antunes et al. 1998; Wu et al. 2000; Carrion and Patterson 2012; Miller et al. 2016; Philippens and Langermans 2021). Several characteristics of this species are highly beneficial for its usage as an animal model, including its small stature and high reproductive capabilities; however, unlike most non-human primates, *C. jacchus* is also characterized by a high frequency of twin-births and hematopoietic chimerism (meaning that non-germline tissue sampled from a single marmoset contains genetic material both from the individual itself as well as from their twin sibling; Hill 1932; Wislocki 1939; Benirschke et al. 1962; Gengozian et al. 1969; Ross et al. 2007; Sweeney et al. 2012; del Rosario et al. 2024) — two biological peculiarities likely to complicate the application of standard population genetic methodologies when conducting genomic studies in this primate.

Although chimerism was initially assumed to be limited to blood samples, Ross et al. (2007) described the presence of chimerism in numerous tissues. This was recently confirmed by del Rosario et al. (2024) in liver, kidney, and brain tissues, though the degrees of chimerism varied, and results continue to suggest that blood samples indeed contain the greatest contribution of sibling nuclei (see Figure 1 of del Rosario et al. 2024, and the accompanying commentary of Chiou and Snyder-Mackler 2024). Moreover, these results also appeared to confirm that chimeric proportions in alternative tissues were proportional to the degree of hematopoietic infiltration (and see Sweeney et al. 2012).

**Figure 1:**
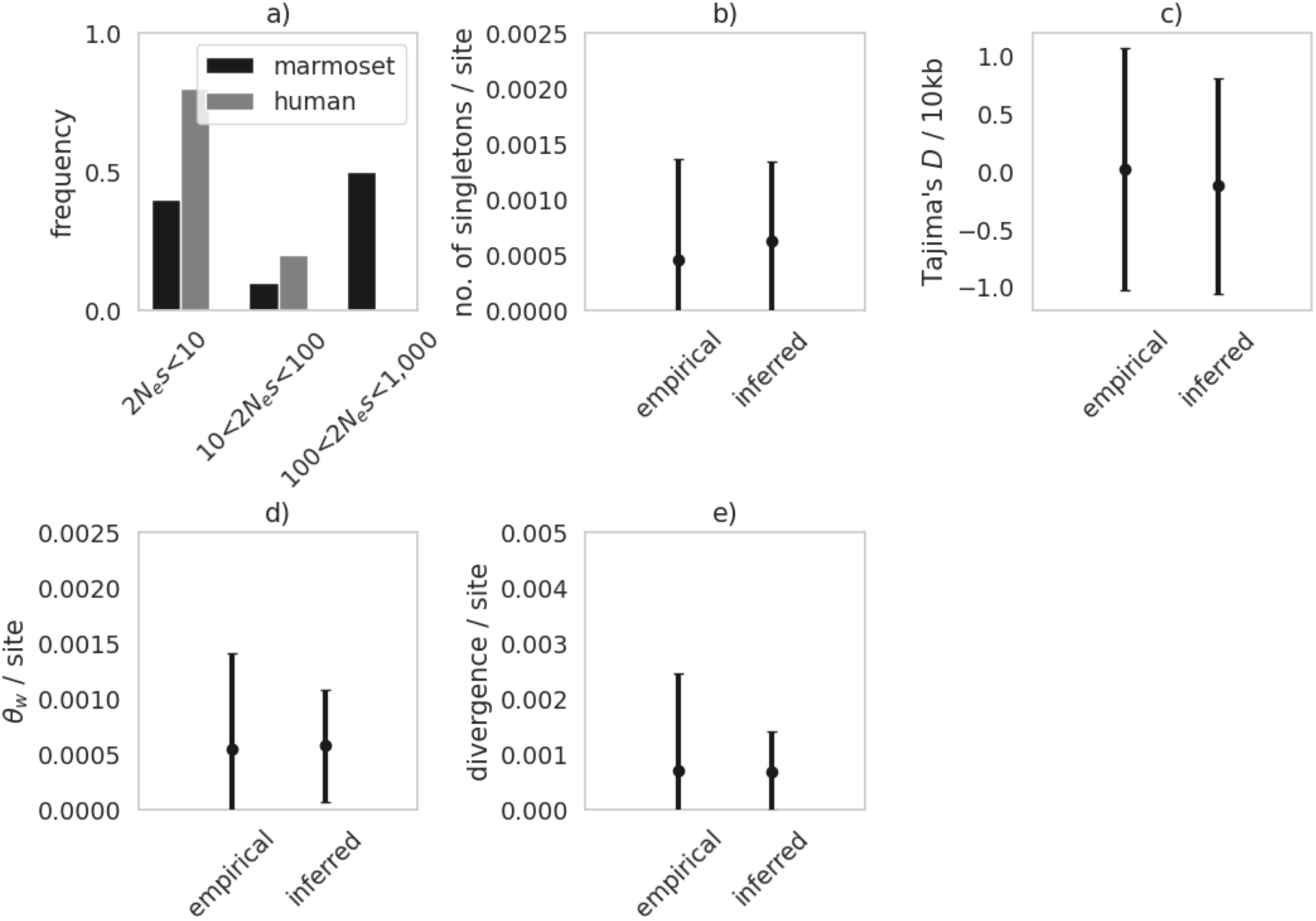
a) Comparison between the best-fitting discrete DFE in the common marmoset inferred in this study (shown in black) and the DFE in humans inferred by Johri et al. 2023 (grey). Exonic mutations were drawn from a DFE comprised of three fixed classes: nearly neutral mutations (i.e., 2*N*_ancestral_ *s* < 10), weakly/moderately deleterious mutations (10 ≤ 2*N*_ancestral_ *s* < 100), and strongly deleterious mutations (100 ≤ 2*N*_ancestral_ *s*). **b-e)** Comparison of four summary statistics—the number of singletons per site, Tajima’s *D* (Tajima 1989) per 10kb, Watterson’s θ_w_ (Watterson 1975) per site, and exonic divergence per site — between the empirical and simulated data under the inferred DFE. Data points represent the mean value, whilst confidence intervals represent standard deviations.

Importantly, this finding implies that it may not be possible to avoid the contributions of chimerism in genetic analyses of the species (e.g., via the use of alternative tissues; Yang et al. 2023). Complicating matters further, circumventing chimeric contributions via the analysis of single births is likely also compromised, as single births are not only rare in common marmosets (Ward et al. 2014) but also tend to themselves be chimeric owing to fetal resorption of a dizygotic twin *in utero* (Jaquish et al. 1996; Windle et al. 1999).

Although the first marmoset genome was published more than a decade ago (The Marmoset Genome Sequencing and Analysis Consortium 2014), population genomic inference in the species thus far has primarily been limited to characterizing genome-wide levels of genetic diversity and divergence (Faulkes et al. 2003; Malukiewicz et al. 2014; Yang et al. 2021; Harris et al. 2023; Yang et al. 2023; Mao et al. 2024) as well as conducting genomic scans based on patterns of genetic hitchhiking (Harris et al. 2014). More recently, Soni et al. (2025d) first investigated the effects of twinning and chimerism on neutral demographic inference, demonstrating a potentially serious mis-inference of population history if neglected, and then designed an approximate Bayesian inference approach accounting for these biological peculiarities to describe a well-fitting history of population size change for this species. Notably, however, no study to date has attempted to similarly model or quantify the effects of twinning and chimerism on the population genetic inference of selection, though the results of Soni et al. (2025d) suggest that this may indeed be important given the role of these biological features in shaping levels and patterns of genomic variation. For example, Harris et al. (2023) found that marmosets had a generally reduced relative heterozygosity compared to other primates, whilst Mao et al. (2024) found that owl monkeys—a closely related, non-chimeric platyrrhine species characterized by frequent singleton births—exhibited a considerably lower divergence to humans compared with marmosets. Yet, because chimerism was unexplored in these studies, it is unclear whether these patterns may be explained by the unusual reproductive dynamic of marmosets alone, or whether non-neutral processes need be invoked.

### Characterizing general selective dynamics: the distribution of fitness effects

The distribution of fitness effects (DFE) describes the spectrum of selection coefficients associated with new mutations, and tends to be relatively stable over long timescales inasmuch as it largely describes the extent of selective constraint. Characterizing the DFE is crucial in evolutionary genomics in that it summarizes the relative proportion of strongly deleterious, weakly deleterious, and neutral mutations, together with the estimated fraction of beneficial variants, in functional genomic regions (see the reviews of Eyre-Walker and Keightley 2007; Keightley and Eyre-Walker 2010; Bank et al. 2014). Since the majority of fitness-altering mutations are deleterious, their ongoing removal through purifying selection—along with associated background selection (BGS) effects (Charlesworth et al. 1993)—continuously shapes genomic diversity. Consequently, accurately quantifying these processes is essential for developing reliable evolutionary baseline models in any given species (Comeron 2014, 2017; Ewing and Jensen 2014, 2016; Johri et al. 2022b; Morales-Arce et al. 2022; Howell et al. 2023; Terbot et al. 2023; Soni and Jensen 2025).

In natural populations, DFE inference approaches utilize polymorphism and/or divergence data to infer either a continuous or discrete distribution of mutational selection coefficients. A commonly used method is the two-step approach of Keightley and Eyre-Walker (2007), in which a demographic model is inferred from synonymous sites in the first step (under the assumption that these sites are evolving neutrally), and then, based on that demographic inference, a DFE is fit to non-synonymous sites (Eyre-Walker and Keightley 2007; Boyko et al. 2008; Eyre-Walker and Keightley 2009; Schneider et al. 2011). More recently, the impact of neglecting BGS effects in such inference was investigated (Johri et al. 2021)—as was the impact of the common neglect of underlying mutation and recombination rate heterogeneity (Soni et al. 2024)—both of which were found to potentially lead to serious mis-inference. Forward-in-time simulation approaches used within an approximate Bayesian setting have thus been developed to jointly infer population history with the DFE, whilst accounting for the effects of selection at linked sites as well as fine-scale mutation and recombination rate heterogeneity (e.g., Johri et al. 2020), utilizing information from the site frequency spectrum (SFS), linkage disequilibrium (LD), as well as divergence data. However, the impact of twinning and chimerism on this inference, of the sort characterizing marmosets, has yet to be explored.

### Characterizing the recent, episodic dynamics of positive and balancing selection

Existing methods for detecting the recent fixation of a beneficial mutation by positive selection rely on the expected changes in patterns of variation at linked sites owing to selective sweep dynamics (see the reviews of Stephan 2019; Charlesworth and Jensen 2021). As would be expected, the nature of these hitchhiking effects will depend on, amongst other factors, the strength and age of the beneficial mutation as well as the local recombination environment. In general, this process is associated with a reduction in local nucleotide diversity (Berry et al. 1991) and a skew in the SFS toward both high-and low-frequency derived alleles within the vicinity of the beneficial fixation (Braverman et al. 1995; Simonsen et al. 1995; Fay and Wu 2000). The theoretical expectations under this model of a single, recent selective sweep have been well described, and provide the theoretical basis for the composite likelihood ratio (CLR) test developed by Kim and Stephan (2002), on which multiple subsequent tests of positive selection have been based. Notably however, it has been demonstrated that alternative evolutionary processes (ranging from genetic drift governed by population history to neutral progeny skew) may closely replicate the patterns associated with a selective sweep (e.g., Jensen et al. 2005; Irwin et al. 2016; Charlesworth and Jensen 2022). One commonly used approach for partially addressing this problem was implemented in the SweepFinder software (Nielsen et al 2005; DeGiorgio et al. 2016), which utilizes a null model derived from the empirically observed SFS as a way of capturing these multi-faceted contributions of alternative processes, and thereby for potentially identifying swept outliers.

Whereas completed selective sweeps reduce variation in their immediate genomic neighborhood, balancing selection maintains genetic variability in populations (see the reviews of Fijarczyk and Babik 2015; Bitarello et al. 2023), potentially over considerable timescales (Lewontin 1987). Based on the genomic signatures left at these varying timescales, Fijarczyk and Babik (2015) characterized different phases of balancing selection, ranging from recent (<0.4*N_e_* generations, where *N_e_* is the effective population size), to intermediate (0.4-4*N_e_* generations), to ancient (>4*N_e_* generations) balancing selection. Notably, the initial trajectory of a newly introduced mutation destined to ultimately experience balancing selection is indistinguishable from that of a partial selective sweep (Soni and Jensen 2024), whereby the newly arisen mutation that has escaped stochastic loss rapidly increases to its equilibrium frequency. These genomic signatures include extended LD due to hitchhiking effects, an excess of intermediate frequency alleles, and a reduction in genetic structure (Schierup et al. 2000; and see the reviews of Crisci et al. 2013; Charlesworth and Jensen 2021). If the selected mutation reaches its equilibrium frequency, it will fluctuate about this frequency if maintained, and the initially generated LD pattern may be broken by subsequent recombination (Wiuf et al. 2004; Charlesworth 2006; Pavlidis et al. 2012). In the case of ancient balancing selection, the targeted allele may continue to segregate as a trans-species polymorphism (Klein et al. 1998; Leffler et al. 2013).

Cheng and DeGiorgio (2020) developed a class of CLR-based methods for the detection of long-term balancing selection, released under the BalLeRMix software package. These methods comprise a mixture model approach, combining the expectation of the SFS under neutrality with the expectation under balancing selection, to infer the expected SFS shape at the putatively selected site and at increasing genomic distances away from that site. As with SweepFinder2, this class of methods utilizes a null model directly derived from the empirical SFS, as opposed to a specified model. Such approaches have been shown to gain power as the balanced mutation segregates over longer timescales (>25*N_e_* generations in age; Soni and Jensen 2024), with new mutations accruing on the balanced haplotype and thereby generating the expected skew in the SFS toward intermediate frequency alleles.

Notably however, as with DFE inference, the impact of the inherent twinning and chimerism characterizing marmosets on expected levels and patterns of genomic variation under models of both positive and balancing selection remains unexplored. Yet, these biological phenomena will represent an essential addition to any appropriate baseline model for this species (Johri et al. 2022a), necessary to comprehensively describe the evolutionary processes certain to occur prior to performing selection scans.

### DFE inference and selection scans in primates

As might be expected, a number of studies have performed DFE inference in humans, beginning with Keightley and Eyre-Walker (2007) who utilized a gene set associated with severe disease or inflammatory response to fit a gamma-distributed DFE. This study estimated a relatively low (∼20%) proportion of effectively neutral mutations and a large proportion (∼40%) of strongly deleterious mutations. A decade later, both Huber et al. (2017) and Johri et al. (2023) inferred a substantially higher proportion of effectively neutral mutations (∼50%) and a smaller proportion of strongly deleterious mutations (∼20%) — differences that can likely at least partially be attributed to differences in the underlying gene sets evaluated. In non-human primates, similar inferences have largely been limited to other genera of the great apes (e.g., Castellano et al. 2019; Tataru and Bataillon 2020), with a notable focus on general regulatory regions (e.g., Simkin et al. 2014; Anderson et al. 2020; Kuderna et al. 2024). More recently, Soni et al. (2025c) performed the first DFE inference in an outgroup to the haplorrhine lineage, the aye-aye (a strepsirrhine), utilizing the annotated chromosome-level genome assembly of Versoza and Pfeifer (2024). Drawing on the two-step framework of Soni and Jensen (2025), the authors inferred a DFE by simulating under the demographic model previously inferred for the species from non-coding regions (sufficiently distant from coding regions to avoid BGS effects) by Terbot et al. (2025). This DFE was found to be characterized by a greater proportion of deleterious variants relative to humans, consistent with the larger inferred long-term effective population size of aye-ayes.

In a similar vein to DFE inference, the majority of genomic scans for positive selection in primates have focused on humans, though numerous studies have been conducted in other great apes (e.g., Enard et al. 2010; Locke et al. 2011; Prüfer et al. 2012; Scally et al. 2012; Bataillon et al. 2015; McManus et al. 2015; Cagan et al. 2016; Munch et al. 2016; Nam et al. 2017; Schmidt et al. 2019) as well as in a handful of biomedically-relevant species (e.g., The Rhesus Macaque Genome Sequencing and Analysis Consortium et al. 2007; Pfeifer 2017b). Recently extending this inference to strepsirrhines, Soni et al. 2025a performed genomic scans for positive and balancing selection in aye-ayes, identifying a number of olfactory-related genes with statistically significant evidence of having experienced long-term balancing selection.

In order to extend this inference to common marmosets as a widely used model system for biomedical research, we have here characterized the effects of twinning and chimerism on general selection inference, and have directly modelled this biology when performing DFE inference and genomic scans, all within the context of the recently estimated population history for this species. In this way, we have robustly identified a number of candidate loci implicated in immune, reproductive, and sensory functions that have strong evidence for having experienced recent or on-going positive and balancing selection effects.

## RESULTS AND DISCUSSION

### Utilizing patterns of exonic divergence to infer the DFE in common marmosets

To quantify fine-scale exonic divergence, we first updated the common marmoset genome included in the 447-way mammalian multiple species alignment (Zoonomia Consortium 2020) to the current reference genome available on NCBI (Yang et al. 2021), before retrieving substitutions along the marmoset branch. Examining the dataset, we found that only a small number of exons exhibited rates of exonic fixation higher than the maximum neutral divergence observed in non-coding regions (Soni et al. 2024b) in 1Mb windows, whereas no exons exceeded the maximum neutral divergence observed in 1kb windows (Supplementary Figure S1) — an anticipated observation given the pervasiveness of purifying selection in functional regions (Charlesworth et al. 1993).

The DFE is expected to remain relatively stable over long evolutionary timescales, and divergence is therefore an informative summary statistic when performing inference of long-term patterns of selection. We utilized forward-in-time simulations in SLiM (Haller and Messer 2023) under the recently published demographic model for the population, and using the modelling framework of Soni et al. (2025d) to account for both twin-births and chimeric sampling, to fit the observed fine-scale patterns of exonic divergence as well as summaries of both genetic variation and the SFS (the number of singletons, Watterson’s θ_w_ [Watterson 1975], and Tajima’s *D* [Tajima 1989]) with a DFE shape consisting of neutral, weakly/moderately deleterious, and strongly deleterious mutational classes. We evaluated this DFE under a divergence time of 0.82 million years (Malukiewicz et al. 2021) between the common marmoset and the closely-related Wied’s black-tufted-ear marmoset (*C. kuhlii*), assuming a generation time of 1.5 years (Okano et al. 2012; Han et al. 2022), by comparing the data summaries between simulated and empirical exonic data in order to quantify a well-fitting DFE. As depicted in Figure 1, a DFE of new mutations characterized by a large proportion of strongly deleterious and nearly neutral variants, and a small proportion of weakly/moderately deleterious mutations, fit the summary statistics observed in common marmosets well. A recent estimate of the DFE from human populations by Johri et al. (2023) has been included for comparison, characterized by a higher density of neutral variants and a considerably lower density of strongly deleterious variants relative to the common marmoset. These patterns are consistent with the much larger long-term effective population size in common marmosets (Soni et al. 2025d), resulting in an increased efficacy of purifying selection (as the strength of selection experienced by an individual mutation is the product of the effective population size, *N_e_*, and the selection coefficient, *s*).

### Evaluating the effects of twinning and chimerism on the inference of recent positive and balancing selection

Prior to performing genomic scans for selection, simulations were utilized to assess whether twinning and chimerism might impact the power and false positive rates (FPR) when employing SFS-based methods for the inference of selective sweeps and balancing selection. Specifically, we performed forward-in-time simulations under an equilibrium population history for both the twin-birth, chimeric-sampling framework outlined in Soni et al. (2025d) and a standard Wright-Fisher (WF) model for comparison, in order to assess statistical performance of the genomic scan approaches implemented in SweepFinder2 (DeGiorgio et al. 2016) and *B_0MAF_* (Cheng and DeGiorgio 2020). Figure 2 provides receiver operating characteristic (ROC) plots across 100 simulated replicates, for two different population-scaled strengths of selection (2*N_e_s* = 100 and 1,000 for sweep inference), and three different times since the introduction of the mutation experiencing negative frequency-dependent selection for balancing selection inference. Given previous results demonstrating that SFS-based methods have little power to detect balanced variants segregating for <25*N* generations (Soni and Jensen 2024), we evaluated introduction times of *τ_b_* = 25*N*, 50*N* and 75*N* generations prior to sampling, where the population size *N* = 10,000. For selective sweep simulations, only replicates in which the beneficial mutation fixed were considered; similarly, for balancing selection simulations, only replicates in which the balanced allele was segregating at the time of sampling were considered. Though twinning and chimerism did not appear to greatly affect sweep inference power, the power to detect balancing selection was notably reduced (Figure 2). As chimerism has been shown to skew the SFS toward intermediate frequency alleles (see Supplementary Figure S6 in Soni et al. 2025d), this likely interferes with the detection of balancing selection owing to the similar signal being produced, whilst the expected selective sweep signal remains rather distinct.

**Figure 2:**
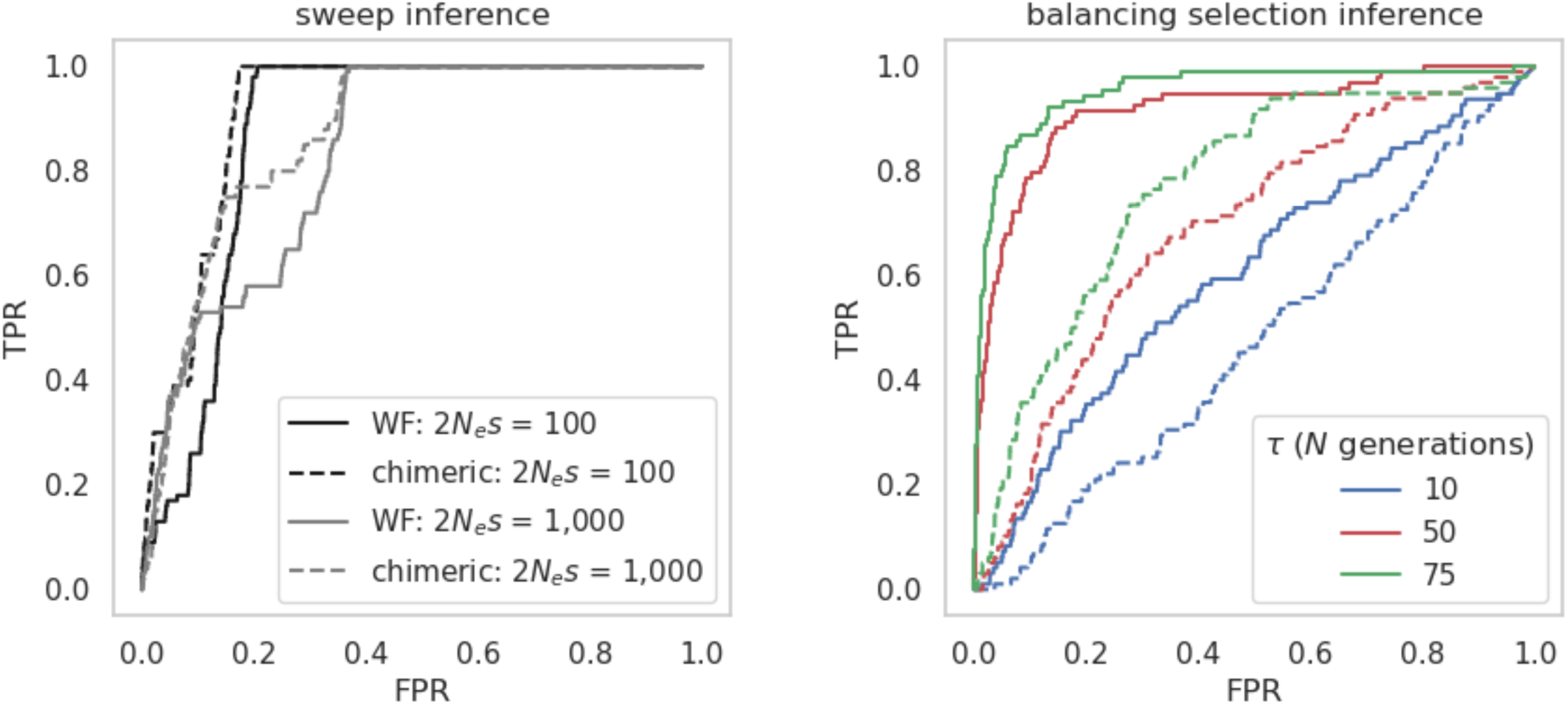
ROC plots based on 100 simulated replicates of a constant population size model for selective sweep (left) and balancing selection (right) inference using the SweepFinder2 and the B*_0MAF_* method, respectively. Solid lines represent standard Wright-Fisher (WF) simulations, whilst dashed lines represent the twin-birth, chimeric-sampling non-WF framework outlined in Soni et al. (2025d), with the false-positive rate (FPR) and the true-positive rate (TPR) provided on the x-axis and y-axis, respectively. For selective sweep inference, power analyses were conducted across two selection regimes — population-scaled strengths of selection of 2*N_e_s* = 100 and 1,000 — with each simulation terminating at the point of fixation of the beneficial mutation. For balancing selection inference, the simulation was terminated at three values of *τ* (the time since the introduction of the balanced mutation): 10*N*, 50*N*, and 75*N* generations, where *N* = 10,000 (for details of simulation and inference schema, see “Materials and Methods”).

Next, we constructed a baseline model consisting of the estimated demographic history for this population (characterized by a population size reduction followed by recovery), twin-births and chimeric sampling (Soni et al. 2025d), the DFE in coding regions capturing purifying and background selection effects (as estimated in this study), as well as mutation and recombination rate heterogeneity by randomly drawing both rates from normal distributions for each 1kb window of our simulated data (for full simulation details, see “Materials and Methods”). For individual selective sweeps, we assessed population-scaled strengths of selection of 2*N_e_s* = 100 and 1,000 at four different times since the introduction of the beneficial mutation (*τ* = 0.2, 0.5, 1, and 2, scaled in *N* generations). As shown in Figure 3, the power to detect selective sweeps is naturally dependent on the strength of selection acting on the beneficial mutation, the time since the introduction of the beneficial mutation, and the window size evaluated. The marmoset demographic model consists of a nearly 70% reduction in population size followed by a partial recovery to roughly half of its ancestral size. Notably, although population bottlenecks can replicate patterns of variation associated with selective sweeps (e.g. Barton 1998; Poh et al. 2014; Harris and Jensen 2020), this reduction in marmosets is not sufficiently severe to strongly reduce power or increase FPRs, and thus considerable power remains for sweep detection in our study. As illustrated in Figure 4, the detection of balancing selection is more sensitive to these factors given the ability of both twinning and chimerism, as well as moderate population bottlenecks, to generate an excess of intermediate frequency alleles. Furthermore, and consistent with previous work (Soni and Jensen 2024), the power to detect balancing selection depends strongly on the time since introduction of the balanced mutation.

**Figure 3:**
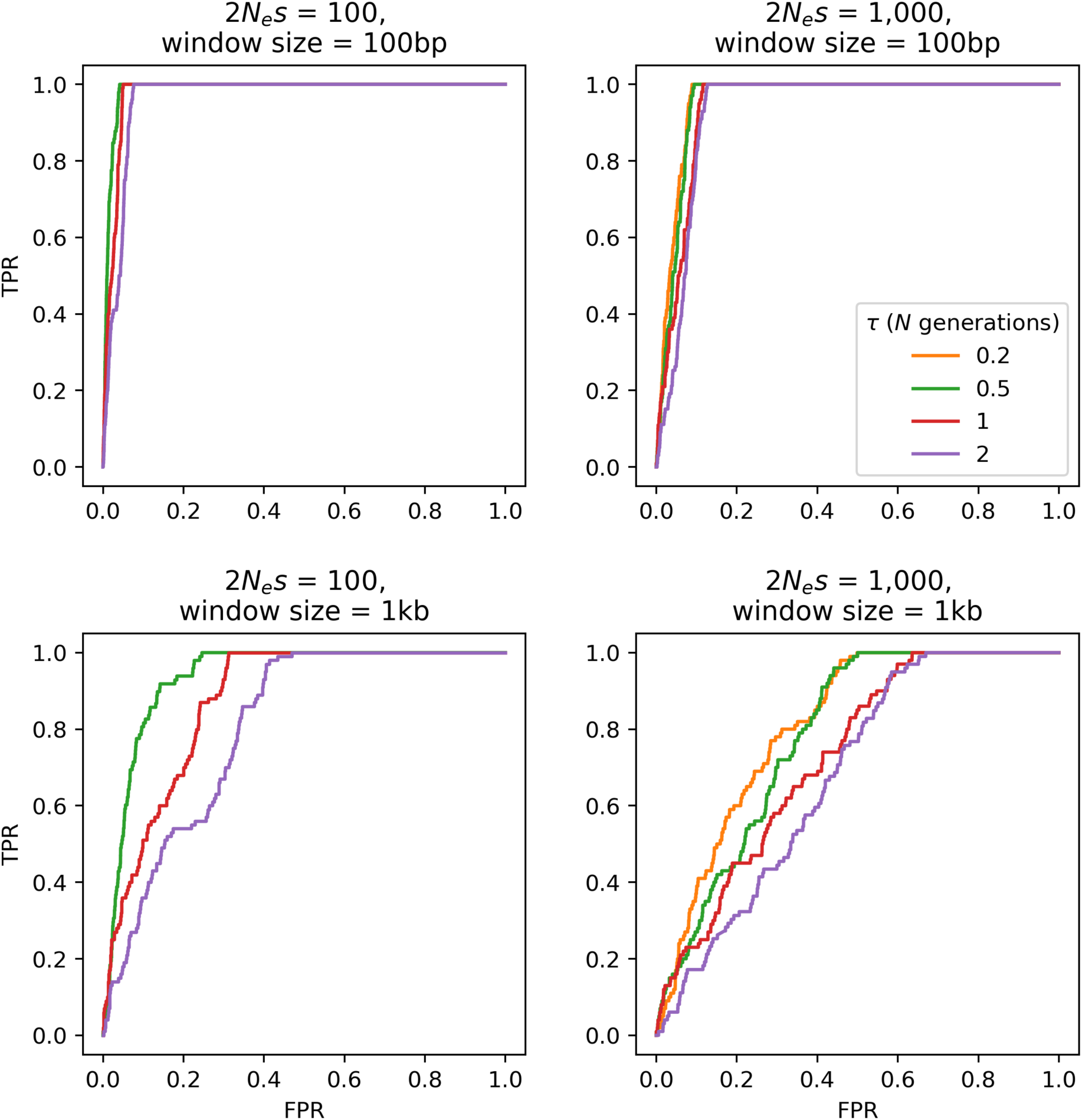
ROC plots based on 100 simulated replicates of the common marmoset demographic model inferred by Soni et al. (2025d) for selective sweep inference using SweepFinder2, with mutation and recombination rates drawn from a normal distribution such that the mean rate per replicate is equal to the fixed rate (see “Materials and Methods” for details). The false-positive rate (FPR) and the true-positive rate (TPR) are provided on the x-axis and y-axis of the ROC plots, respectively. Power analyses were conducted across two selection regimes—population-scaled strengths of selection of 2*N_e_s* = 100 and 1,000 — four times of introduction of the beneficial mutation (*τ* = 0.2*N*, 0.5*N*, 1*N*, 2*N*; where *N* = 61,898), and two window sizes (100 bp and 1 kb). Note that no ROC could be plotted for simulation schemata in which none of the beneficial mutations reached fixation at the time of sampling.

**Figure 4:**
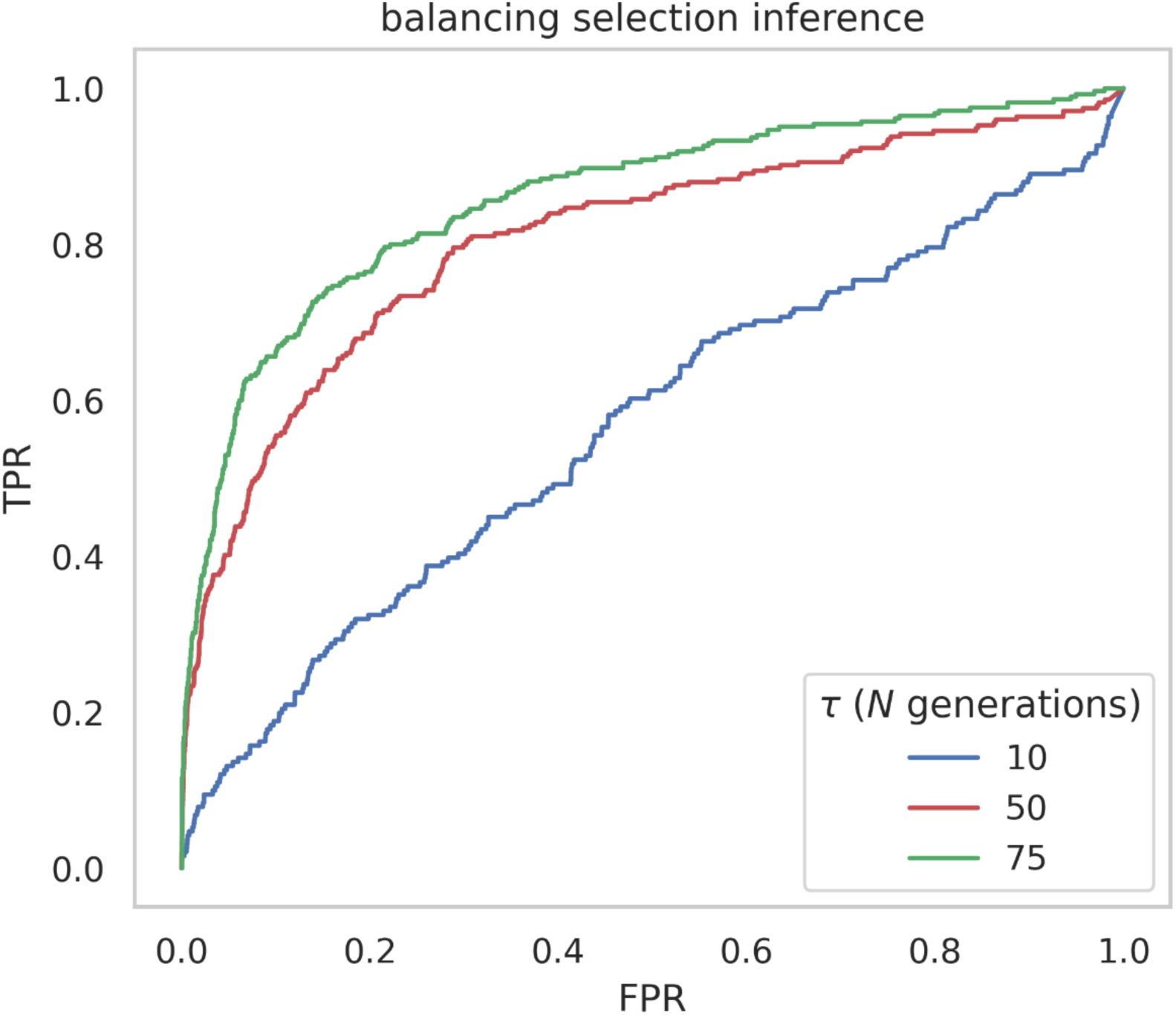
ROC plots based on 100 simulated replicates of the common marmoset demo-graphic model inferred by Soni et al. (2025d) for balancing selection inference using *B_0MAF_*, with mutation and recombination rates drawn from a uniform distribution such that the mean rate per replicate is equal to the fixed rate (see “Materials and Methods” for details). The false-positive rate (FPR) and the true-positive rate (TPR) are provided on the x-axis and y-axis of the ROC plots, respectively. Power analyses were conducted across three times of introduction of the balanced mutation (*τ* = 10*N*, 50*N,* and 75*N* generations; where *N* = 61,898).

Based upon these results, and in order to robustly identify candidate loci experiencing recent positive and/or balancing selection within the context of this marmoset biology and population history, we performed genome-wide selection scans using the CLR methods SweepFinder2 (DeGiorgio et al. 2016) to detect signals of positive selection (performing sweep inference at each SNP individually) and *B_0MAF_* (Cheng and DeGiorgio 2020) to detect signals of balancing selection (performing sweep inference in 1 kb windows with a 50 bp step size). Although outlier approaches — i.e., approaches that assume that genes in the chosen tail of the distribution (often 5% or 1%) are likely sweep candidates, regardless of the underlying evolutionary model (Harris et al. 2018, and see discussions in Howell et al. 2023; Jensen 2023; Johri et al. 2023; Terbot et al. 2023) — are commonly used to identify candidate regions experiencing positive selection, such approaches have been shown to be associated with high false-positive rates (Teshima et al. 2006; Thornton and Jensen 2007; Jensen et al. 2008; Jensen 2023; Soni et al. 2023). We therefore instead constructed an evolutionarily appropriate baseline model accounting for commonly operating evolutionary processes, as recommended by Johri et al. (2022b, and see Figures 5 and 6 in Johri et al. 2022a for diagrams summarizing important considerations for constructing such genomic baseline models). Based on the maximum CLR values observed under neutrality across simulation replicates of the inferred marmoset population history with twinning and chimerism, we set the null thresholds for positive and balancing selection inference as 82.16 for SweepFinder2 and 87.57 for *B_0MAF_*, respectively (see “Materials and Methods” for details), and identified candidate regions as any empirical CLR values in excess of these thresholds.

**Figure 5:**
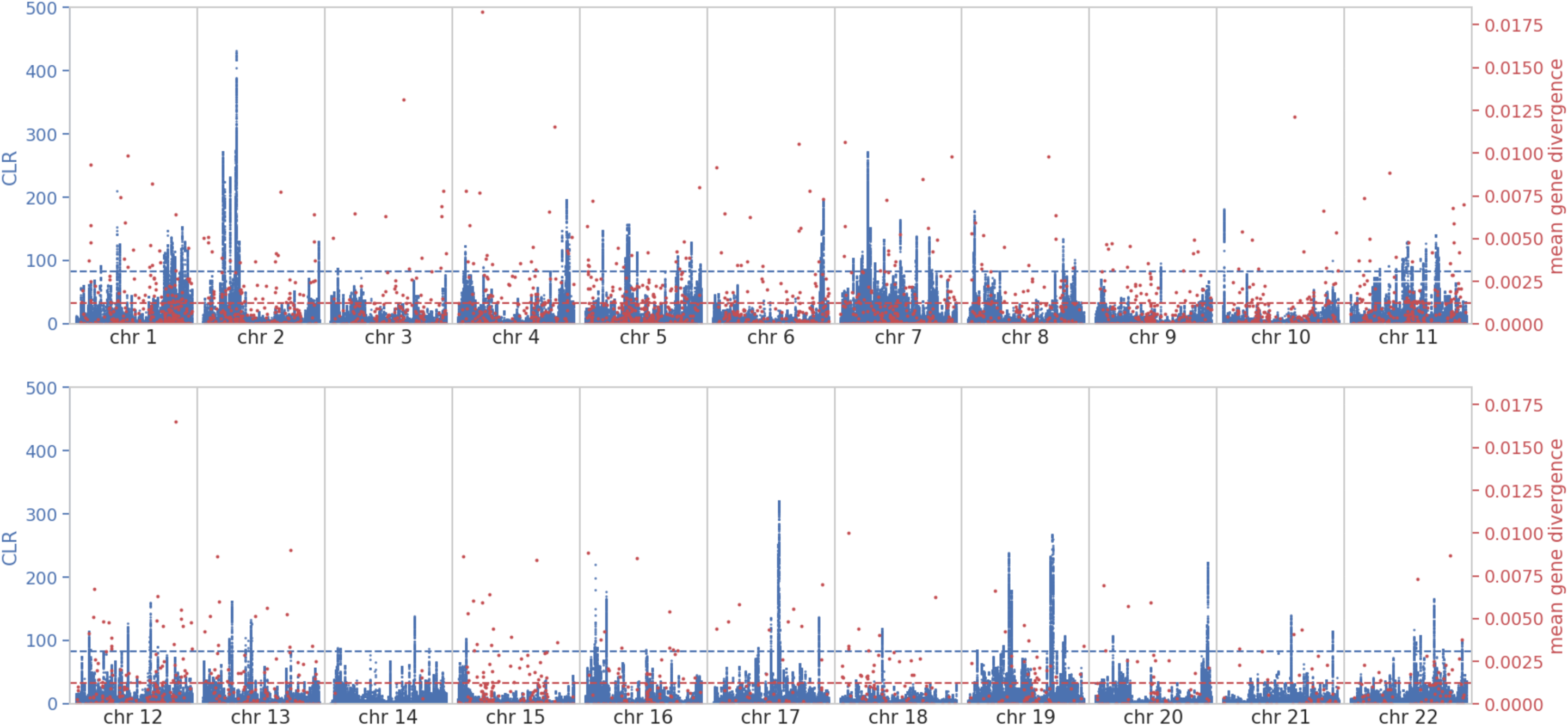
Genome-wide selective sweep scan results using SweepFinder2 (shown in blue) and empirical exonic divergence (red). The x-axis shows the position along each autosome (chromosomes 1-22), the left y-axis shows the composite likelihood ratio (CLR) value of the sweep statistic at each SNP, and the right y-axis shows the mean gene divergence at each SNP. The horizontal blue dashed line represents the null threshold for sweep detection, and the horizontal red dashed line represents the 75^th^ percentile neutral divergence.

**Figure 6:**
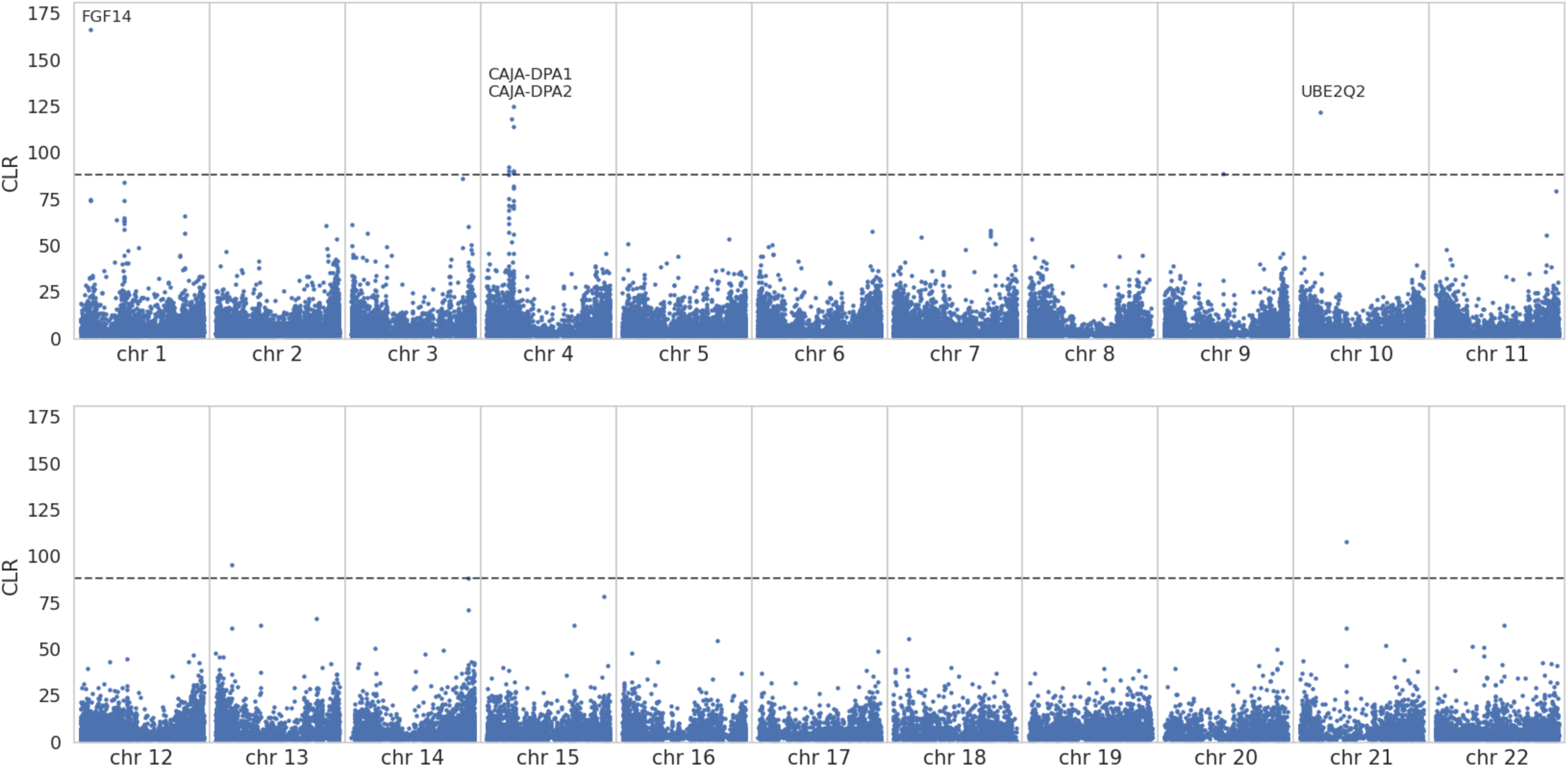
Genome-wide balancing selection scan results using *B_0MAF_*. The x-axis shows the position along each autosome (chromosomes 1-22) and the y-axis shows the composite likelihood ratio (CLR) value of the balancing selection statistic at each SNP. The horizontal dashed line represents the null threshold for detection.

### Signatures of positive selection in the common marmoset genome

Utilizing the thresholds from the evolutionary baseline model, we identified 10,599 loci with statistically significant evidence of having experienced selective sweep effects, mapping to 216 genes within the marmoset genome (see Figure 5 for the genome-wide scan results for SweepFinder2, Supplementary Figures S2-S23 for scan results along individual chromosomes, and Supplementary File S1 for candidate gene regions with CLR values greater than the null threshold). Performing a gene functional analysis of these 216 candidates using the Database for Annotation, Visualization, and Integrated Discovery (DAVID; Ma et al. 2023), we identified several categories related to cellular components, sequence features, and gene regulation (Supplementary Table S1). Additionally, we compared the identified highly divergent genes on the *C. jacchus* branch in the multi-species alignment with these sweep candidates (see Supplementary Figures S2-S23 for per-chromosome exonic divergence overlayed with the genomic scan results). These analyses highlighted a number of genes and categories of interest:

i. Immune-related genes are among the most rapidly evolving across vertebrates, due to the evolutionary arms race in response to pathogen exposure (e.g., George et al. 2011; Rausell and Telenti 2014; Soni et al. 2025c). Three tumor-related genes were identified as candidates for recent positive selection in the peaks of the likelihood surface. Specifically, loci in the gene *EPHB1* had the highest CLR value in our genome scans—a gene that plays a critical role in immune cell development and function, and that has been implicated in several nervous system diseases, cardiovascular diseases, and cancers (Xie et al. 2024). The tumor suppressor *RPH3AL* was both a candidate for recent positive selection and exhibited an unusually high rate of fixations. The *BRAF* gene regulating cell growth is a proto-oncogene in which strongly activating mutations have been argued to experience selective sweep dynamics in humans (Gopal et al. 2019). Additionally, *TSPAN2* and *TSBP1* exhibited a rate of divergence higher than the 99^th^ percentile neutral divergence (as estimated by Soni et al. 2025b). *TSPAN2* plays a role in cell motility and has been implicated in both nervous system development and cancer progression (Otsubo et al. 2014, and see the review of Yaseen et al. 2017). The most highly diverged gene identified in our dataset, *TSBP1,* is located in the major-histocompatibility complex (MHC). Hoh et al. (2020) identified a genomic region encompassing *TSBP1* as undergoing positive selection in native human populations in North Borneo, hypothesizing that the *Plasmodium* parasite endemic to the region was the driver of local adaptation. Given that the common marmoset is one of numerous New World monkeys that are targets of infection by *P. brasilianum* (Alvarenga et al. 2017), this parasite may similarly be driving these dynamics.
ii. The candidate gene *CDC14B* is also noteworthy in that its retrogene *CDC14B2* originated by retroduplication in the hominoid ancestor ∼18-25 million years ago (mya; Marques et al. 2008), with evidence of experiencing positive selection in African apes ∼7-12 mya (Rosso et al. 2008). In mice, *CDC14B2*-deficient cells have been shown to accumulate more endogenous DNA damage than wild-type cells, consistent with premature aging (Wei et al. 2011)—a result of particular interest given that marmosets remain as a widely-used model organism for the study of neurodegeneration and aging (Perez-Cruz and Rodriguez-Callejas 2023).
iii. Given that many of the unique aspects of marmoset biology revolve around reproduction, identifying genes involved in ovulation and gestation is of considerable biomedical interest. Three genes linked to reproductive function were observed to fall in the high-divergence gene set. *SPACA7*, for example, plays a vital role in spermatogenesis (Aisha and Yenugu 2023), and has been argued to be experiencing positive selection across the primate clade (van der Lee et al. 2017); additionally, there is evidence that *ZP2,* involved in female fertilization and the formation of the mammalian egg coat, has experienced positive selection across the mammalian clade (Swanson et al. 2001).
iv. A strongly enriched functional category arising from the joint candidates were genes related to calcium. As marmosets are characterized by a lack of readily available calcium in their diet and a poor ability to digest this mineral (Jarcho et al. 2013) — indeed, marmosets, and in particular lactating females, have been shown to exhibit a preference for calcium lactate solutions over plain water (Power et al. 1999) — this result may imply on-going selective pressures related to this mineral intake.
v. Marmosets, who live in extended social family groups and engage in cooperative breeding, exhibit a rich set of vocalizations to identify and communicate information about predators and food, as well as to convey biologically important information such as identity, sex, and emotional states (Seyfarth and Cheney 2003; Agamaite et al. 2015; Lamothe et al. 2025). Notably in this regard, the gene *TMEM145*, which plays an important role in hearing, specifically in the structure and function of outer hair cell stereocilia in the inner ear (Roh et al. 2025), was both highly divergent (>75^th^ percentile of neutral divergence) and a selective sweep candidate.

### Signatures of balancing selection in the common marmoset genome

By contrast, only 14 strongly supported candidate regions met our null threshold for balancing selection inference, mapping to four genes within the common marmoset genome. Figure 6 provides the genome-wide scan results for *B_0MAF_* (and see Supplementary Figures S24-S45 for the scan results along individual chromosomes, and Supplementary File S1 for candidate windows exhibiting CLR values greater than the null threshold). We were unable to perform a gene functional analysis with such a small number of candidate genes, and manually curated this set as an alternative, identifying two MHC genes: *CAJA-DPA1* and *CAJA-DPA2*. MHC loci have frequently been implicated in balancing selection in multiple species (see the review by Radwan et al. 2020), and the sharing of polymorphism is a signature of balancing selection that has been identified both between human populations (Soni et al. 2022) and across apes (Leffler et al. 2013; Teixeira et al. 2015). Additional support for balancing selection in-*DPA1* genes was recently argued in the form of trans-species polymorphisms shared among the African apes (Fortier and Pritchard 2025). Notably, Antunes et al. (1998) found that the *CAJA-DP* region may be altered in appearance in common marmosets, given that other higher primate species have three distinct functional MHC class II regions (-*DR*,-*DQ*, and-*DP*), whilst *CAJA-DPA1* could only be detected in low quantities of PCR product in the common marmoset, suggesting that-DR and-DQ represent the main functional MHC class II regions in this species.

## CONCLUDING THOUGHTS

In this study we have inferred both recent and long-term patterns of natural selection in the common marmoset genome. We have estimated a well-fitting DFE to model the effects of purifying and background selection, which additionally accounts for observed patterns of exonic divergence. We found evidence of an increased proportion of newly arising strongly deleterious variants in marmosets relative to humans, potentially related to their larger estimated long-term effective population size.

Additionally, in order to perform the first large-scale scans for loci having experienced selective sweeps and/or balancing selection in the marmoset genome, we generated an evolutionary baseline model utilizing the recently estimated population history of Soni et al. (2025d), our inferred DFE, as well as existing knowledge regarding mutation and recombination rate heterogeneity, taking into account the reproductive dynamics of common twin-births and chimeric sampling particular to this species in this model construction. Utilizing this conservative approach to reduce false-positive rates, a number of genes with relevance to both biomedical and evolutionary interest were identified, including genes related to immune, sensory-related, and reproductive functions. Notably, these gene sets particularly highlighted MHC genes as having strong evidence for experiencing long-term balancing selection, consistent with an accumulating body of work across the primate clade.

## MATERIALS AND METHODS

### Animal subjects

Animals previously housed at the New England Primate Research Center were maintained in accordance with the guidelines of the Harvard Medical School Standing Committee on Animals and the Guide for Care and Use of Laboratory Animals of the Institute of Laboratory Animal Resources, National Research Council. All samples were collected during routine veterinary care under approved protocols.

### Whole genome, population-level data

We utilized previously collected blood samples from 15 common marmosets to extract DNA using the FlexiGene kit (Qiagen, Valencia, CA) following the standard protocol without any modifications. For each sample, we prepared a PCR-free library that was sequenced to a target coverage of 35X on a DNBseq platform at the Beijing Genomics Institute, resulting in 150bp paired-end reads. We pre-processed the raw reads following standard quality-control practices (Pfeifer 2017a) using SOAPnuke v.1.5.6, trimming adapters and removing both low-quality and polyX tails (with the parameters: “*-n* 0.01 *-l* 20 *-q* 0.3 *-A* 0.25 *--cutAdaptor-Q* 2 *-G--polyX--minLen* 150”; Chen et al. 2018). We mapped the pre-processed reads with BWA-MEM v.0.7.17 (Li and Durbin 2009) to the common marmoset reference genome (mCalJa1.2.pat.X; GenBank accession number: GCA_011100555.2; Yang et al. 2021). From these read mappings, we identified and marked duplicates using Picard’s *MarkDuplicates* embedded within the Genome Analysis Toolkit (GATK) v.4.2.6.1 (van der Auwera and O’Connor 2020). Following the recommendations of the developers, we generated a gVCF for each sample using the GATK *HaplotypeCaller* (providing the “*--pcr-indel-model* NONE” flag to account for the fact that the reads originated from PCR-free libraries), combined gVCFs across samples with *CombineGVCFs*, and jointly genotyped them using *GenotypeGVCFs.* In all steps, we included both variant and invariant sites (using the “*--emit-ref-confidence* BP_RESOLUTION” and “*--include-non-variant-sites*” flags in the *HaplotypeCaller* and *GenotypeGVCFs*, respectively) genotyped in all individuals (“AN = 30”). To filter out spurious sites, we applied GATK’s hard filter criteria (i.e., *FS* > 60.0; *MQ* < 40.0; *MQRankSum* <-12.5; *QD* < 2.0; *QUAL* < 30.0; *ReadPosRankSum* <-8.0; *SOR* > 3.0; with acronyms as defined by GATK) as well as an additional filter criterion based on read coverage (0.5 × *DP_ind_* ≤ *DP_ind_* ≤ 2.0 × *DP_ind_*), and removed any sites located within repetitive regions prone to mapping errors with short-read data. Lastly, we limited the dataset to autosomes (chromosomes 1-22) for downstream analyses (Supplementary Table S2).

### Calculating exonic divergence

We updated the common marmoset genome included in the 447-way multiple species alignment (Zoonomia Consortium 2020) to the current reference genome available on NCBI (GenBank accession number: GCA_011100555.2; Yang et al. 2021) by first removing the included marmoset genome using the HAL v.2.2 (Hickey et al. 2013) *halRemoveGenome* function and then extracting the neighboring sequences (i.e., the genomes of the closely-related Wied’s black-tufted-ear marmoset, *C. kuhlii*, and the ancestral PrimateAnc232) using *hal2fasta*. Afterward, we realigned these genomes with the current marmoset genome assembly in Cactus v.2.9.2 (Armstrong et al. 2020), preserving the original branch lengths. Finally, we reintegrated the updated subalignment using HAL’s *halReplaceGenome*.

To estimate fine-scale exonic divergence, we used HAL’s *halSummarizeMutations* function to identify fixed differences in exons along the marmoset lineage (i.e., between the genomes of the common marmoset and the ancestral PrimateAnc232). To mitigate the effects of fragmentation present in the *C. kuhlii* genome — which, unlike the common marmoset genome, is currently at the scaffold-level — we kept only those exons found in alignments longer than 10 kb in length. Lastly, we masked any point mutations known to be polymorphic in the species, and calculated the rate of exonic divergence by dividing the number of substitutions by the number of accessible sites in each exon.

### DFE inference

In order to fit a DFE to the coding regions of the common marmoset genome, we used forward-in-time simulations in SLiM v.4.23 (Haller and Messer 2023) to simulate 100 exonic regions, each corresponding to the mean length of exons greater than 1 kb observed in the empirical data (i.e., 3,209 bp). More specifically, we performed simulations under the recently published demographic model for the species using the modelling framework of Soni et al. (2025d) to account for both twin-births and chimeric sampling, and including a 10*N_ancestral_* generation burn-in time prior to the start of the demographic model (where *N*_ancestral_ is the initial population size of 61,198 individuals). We assumed a branch split time of 0.82 million years (Malukiewicz et al. 2021) and a generation time of 1.5 years (Okano et al. 2012; Han et al. 2022), and randomly drew mutation and recombination rates for each replicate from a normal distribution, such that the mean rates across all 100 simulation replicates were equal to the mean rates inferred in closely related primates (using a mean mutation rate of 0.81 × 10^-8^ per base pair per generation and a mean recombination rate of 1cM/Mb, as per Soni et al. 2025d). Following Johri et al. (2020), we drew exonic mutations from a DFE comprised of three fixed classes: nearly neutral mutations (2*N*_ancestral_ *s* < 10, where *N_ancestral_* is the ancestral population size and *s* is the reduction in fitness of the mutant homozygote relative to wild-type), weakly/moderately deleterious mutations (10 ≤ 2*N*_ancestral_ *s* < 100), and strongly deleterious mutations (100 ≤ 2*N*_ancestral_ *s*).

To infer the DFE, we then performed a grid search by varying the fractions of mutations obtained from each category. Specifically, for each parameter combination, we simulated 100 replicates using the DFE of human functional regions as a starting point (see Johri et al. 2023 for details), and compared the fit of four summary statistics between our empirical and simulated data: Watterson’s θ_w_ (Watterson 1975) per site, Tajima’s *D* (Tajima 1989) per 10kb, the number of singletons per site, and exonic divergence per site. We used pylibseq v.1.8.3 (Thornton 2003) to calculate Watterson’s θ_w_, Tajima’s *D*, and the number of singletons across 10 kb windows with a 5 kb step size, whereas exonic divergence was calculated as the number of fixations that occurred in our simulated population post-burn-in to enable a direct comparison to the empirically observed number of substitutions along the marmoset branch.

### Evaluating the effects of chimerism on genome scans

In order to evaluate the effects of chimerism on genome scans for positive and balancing selection, we simulated 100 replicates of a single marmoset population using SLiM v.4.3 (Haller and Messer 2023). To mimic the genomic architecture of the species, we simulated three functional regions — comprised of nine 130 bp exons separated by 1,591 bp introns — isolated by 16,489 bp intergenic regions, for a total region length of 91,161 bp in each replicate. We modelled mutations in intronic and intergenic regions as neutral, and drew exonic mutations from a DFE comprised of four fixed classes with frequencies denoted by *f_i_*: effectively neutral mutations *f_0_* (i.e., 0 ≤ 2*N_ancestral_s* < 1), weakly deleterious mutations *f_1_* (1 ≤ 2*N_e_s* < 10), moderately deleterious mutations *f_2_* (10 ≤ 2*N_ancestral_s* < 100), and strongly deleterious mutations *f_3_* with (100 ≤ 2*N_ancestral_s* < 2*N_e_*), with *s* drawn from a uniform distribution within each bin. We implemented two modelling schemes: an equilibrium population history, and the inferred marmoset demographic model of Soni et al. (2025d). For the equilibrium model, we simulated under the twin-birth, chimeric-sampling non-WF framework outlined in Soni et al. (2025d) and a WF model for comparison, both using a 10*N_ancestral_* burn-in time (where *N_ancestral_* = 10,000). For the marmoset demographic model, we simulated under the twin-birth, chimeric-sampling non-WF framework only, again with a 10*N_ancestral_* burn-in time (where *N_ancestral_* = 61,898). In all cases, we sampled 15 individuals at the end of each simulation. In each simulation replicate, a single positively selected mutation was introduced. For the selective sweep analysis, we simulated two beneficial population-scaled selection coefficients — 2*N_e_s* = [100, 1,000] — and sampled the population at *τ* = [0.1, 0.2, 0.5], where *τ* is the time since fixation of the beneficial mutation in *N* generations in the equilibrium model; for the demographic model simulations, we introduced beneficial mutations at times [0.2, 0.5, 1, 2] *N_ancestral_* generations prior to the end of the simulation. For the balancing selection analysis, we introduced the balanced mutation at *τ_b_* = [10, 50, 75], where *τ_b_* is the time since the introduction of the balanced mutation in *N* generations.

We modelled the balanced mutation to experience negative frequency-dependent selection, i.e., in a manner such that the selection coefficient was dependent on the frequency in the population: *S_bp_* = *F_eq_* – *F_bp_*, where *S_bp_* is the selection coefficient of the balanced mutation, *F_eq_* is the equilibrium frequency of the balanced mutation (here set to 0.5), and *F_bp_* is the frequency of the balanced mutation in the population. We implemented all simulations such that if the selective sweep failed to fix, or if the balanced mutation was fixed or lost from the population, the simulation would restart at the point of introduction of the selected mutation.

With these simulations on hand, we then performed selective sweep inference using SweepFinder2 v.1.0. (DeGiorgio et al. 2016) to detect signals of positive selection (using the command: “*SweepFinder2 –lu* GridFile FreqFile SpectFile OutFile”) and *B_0MAF_* (Cheng and DeGiorgio 2020) to detect signals of balancing selection (using the command: “*python3 BalLeRMix+_v1.py-I* FreqFile *--spect* SfsFile *-o* OutFile *–noSub – MAF –rec* 1e-8”), with analyses limited to folded allele frequencies and polymorphic sites only, performing inference at each SNP. Under the demographic model, balancing selection inference was performed in 1kb windows with a 50bp step size, to mimic the conditions of the empirical inference performed in this study.

### Generating null thresholds for selection inference

In order to generate null thresholds for the inference of selection, we simulated the demographic model of the species (based on the twin-birth, chimeric-sampling non-WF framework outlined in Soni et al. 2025d), consisting of a relatively recent population bottleneck followed by a recovery via exponential growth to roughly half the ancestral size. In brief, based on this demographic history, we simulated 10 replicates for each of the 22 autosomes in the marmoset genome in SLiM v.4.3 (Haller and Messer 2023), simulating only neutral regions that were at least 10 kb from the nearest coding region, in order to avoid the biasing effects of purifying and background selection. Following Soni et al. (2025d), we assumed a mutation rate of 0.81 × 10^-8^ per base pair per generation and a recombination rate of 1cM/Mb. To reduce FPRs, we conservatively used the maximum CLR value observed across all null model simulations (i.e., the highest value observed in the absence of positive or balancing selection) as the null threshold for the selection scans with SweepFinder2 v.1.0 (DeGiorgio et al. 2016) and *B_0MAF_* (Cheng and DeGiorgio 2020) on the empirical data.

### Inferring recent positive and balancing selection in the marmoset genome

With the null thresholds, we applied the inference schema discussed above with SweepFinder2 and *B_0MAF_* on the empirical autosomal data. We performed sweep inference at each SNP and balancing selection inference across 1 kb windows, with a step size of 50 bp. We manually curated our candidate loci by identifying genes under the significant likelihood surface, and evaluated the obtained dataset via the NCBI database (Sayers et al. 2022) and Expression Atlas (Madeira et al. 2022) in order to identify function and expression patterns in different primate species. Additionally, we used the Database for Annotation, Visualization, and Integrated Discovery (DAVID; Ma et al. 2023) to perform a Gene Ontology analysis (The Gene Ontology Consortium 2023).

### Identifying synonymous and non-synonymous mutations

Extensive genome fragmentation can impact comparative genomic analyses. As many primate genome assemblies present in the 447-way multiple species alignment are currently still at the scaffold (rather than chromosome) level, we limited our analyses to a subset of high-quality genomes representing different branches along the primate clade, with humans (hg38; Schneider et al. 2017) as a representative for the haplorrhines, common marmosets (mCalJa1.2.pat.X; Yang et al. 2021) as a representative for the platyrrhines, and aye-ayes (DMad_hybrid; Versoza and Pfeifer 2024) as a representative for strepsirrhines. Additionally, we further curated the alignments to only include exonic regions of the genes identified by SweepFinder2 (*n* = 35) using the genome annotation of the common marmoset assembly (Yang et al. 2021) and the *hal2maf* function implemented in Cactus v.2.9.3 (Armstrong et al. 2020). Using a custom script, we identified fixed differences between marmosets and both humans and aye-ayes (i.e., positions at which both humans and aye-ayes carry the same nucleotide but marmosets carry a different nucleotide), removing any sites known to segregate in the species. We then utilized SnpEff v.5.2.1 (Cingolani et al. 2012) to identify synonymous (i.e., *SYNONYMOUS_CODING*, *SYNONYMOUS_START*, and *SYNONYMOUS_STOP*) and non-synonymous (i.e., *NON_SYNONYMOUS_CODING*, *NON_SYNONYMOUS_START*, and *NON_SYNONYMOUS_STOP*) mutations.

### Identifying highly divergent genes

We calculated mean divergence per gene, and performed gene functional analysis using DAVID (Ma et al. 2023) on the subset of genes that overlapped between sweep candidates, and those with a mean divergence value greater than the 75^th^, 95^th^, and 99^th^ percentiles of neutral divergence.

## ACKNOWLEDGEMENTS

We would like to thank Eric J. Vallender for providing the marmoset samples used in this study, and the members of the Jensen and Pfeifer Labs for helpful comments and discussions. Computations were performed on the Sol supercomputer at Arizona State University (Jennewein et al. 2023).

