## Supplementary Materials for "Investigating the effects of chimerism on the inference of selection: quantifying genomic targets of purifying, positive, and balancing selection in common marmosets (*Callithrix jacchus*)"

**Supplementary Table S1**

| Analysis | Category | Term | *p*-value | Fold Enrichment | FDR |
|---|---|---|---|---|---|
| All selective sweep candidates | UP_SEQ_FEATURE | COMPBIAS:Basic and acidic residues | 3.09E-06 | 1.483 | 0.003 |
| | UP_SEQ_FEATURE | REGION:Disordered | 6.54E-05 | 1.185 | 0.035 |
| | UP_SEQ_FEATURE | COMPBIAS:Polar residues | 1.33E-04 | 1.429 | 0.038 |
| | UP_SEQ_FEATURE | COMPBIAS:Basic residues | 1.44E-04 | 1.778 | 0.038 |
| | UP_KW_PTM | KW-0597~Phosphoprotein | 2.96E-04 | 1.234 | 0.006 |
| | UP_KW_PTM | KW-0007~Acetylation | 5.64E-04 | 1.507 | 0.006 |
| | UP_KW_CELLULAR_COMPONENT | KW-0333~Golgi apparatus | 9.12E-04 | 2.164 | 0.035 |
| Peak selective sweep genes | UP_KW_CELLULAR_COMPONENT | KW-0539~Nucleus | 1.97E-03 | 1.870 | 0.034 |
| Peak selective sweep genes with high mean divergence | UP_SEQ_FEATURE | CARBOHYD:N-linked (GlcNAc...) asparagine | 5.53E-07 | 1.373 | 0.002 |
| | GOTERM_CC_DIRECT | GO:0062023~collagen-containing extracellular matrix | 2.66E-05 | 2.392 | 0.007 |
| | GOTERM_CC_DIRECT | GO:0016020~membrane | 3.84E-05 | 1.265 | 0.007 |
| | GOTERM_CC_DIRECT | GO:0005886~plasma membrane | 4.29E-05 | 1.257 | 0.007 |
| | UP_KW_LIGAND | KW-0106~Calcium | 1.16E-04 | 1.627 | 0.004 |
| | GOTERM_CC_DIRECT | GO:0005576~extracellular region | 1.33E-04 | 1.427 | 0.017 |
| | UP_KW_PTM | KW-0325~Glycoprotein | 2.24E-04 | 1.221 | 0.006 |
| | UP_KW_CELLULAR_COMPONENT | KW-0966~Cell projection | 8.31E-04 | 1.491 | 0.027 |

**Supplementary Table S1:** Enriched gene categories from functional analysis with DAVID, with false discovery rates (FDR) and *p*-values <0.05 for all selective sweep candidate genes (top), genes at likelihood surface peaks (middle), and genes that have both high divergence and are at likelihood surface peaks (bottom).

**Supplementary Table S2**

| chrom | length | # SNPs | # invariant sites | Ts/Tv |
|---|---|---|---|---|
| 1 | 216,975,769 | 257,119 | 47,237,148 | 2.20 |
| 2 | 202,808,461 | 222,710 | 40,777,876 | 2.11 |
| 3 | 189,455,076 | 162,252 | 28,397,479 | 2.01 |
| 4 | 173,414,057 | 228,055 | 37,833,102 | 2.04 |
| 5 | 161,716,902 | 213,640 | 51,551,915 | 2.16 |
| 6 | 159,674,559 | 205,517 | 33,317,930 | 2.13 |
| 7 | 156,129,252 | 237,836 | 43,121,041 | 2.18 |
| 8 | 126,104,592 | 169,719 | 26,637,950 | 2.10 |
| 9 | 132,900,640 | 152,330 | 29,901,412 | 2.10 |
| 10 | 136,971,485 | 201,931 | 33,828,189 | 2.10 |
| 11 | 128,529,401 | 178,097 | 31,238,359 | 2.13 |
| 12 | 123,697,827 | 188,874 | 33,767,759 | 2.25 |
| 13 | 117,638,787 | 142,340 | 29,270,661 | 2.11 |
| 14 | 112,969,529 | 149,207 | 27,988,452 | 2.09 |
| 15 | 98,621,277 | 145,110 | 24,183,383 | 2.11 |
| 16 | 98,110,366 | 112,237 | 19,037,643 | 1.99 |
| 17 | 74,700,100 | 88,568 | 14,898,892 | 2.03 |
| 18 | 47,063,576 | 70,276 | 12,475,879 | 2.05 |
| 19 | 50,540,917 | 111,593 | 15,956,698 | 2.11 |
| 20 | 44,412,365 | 76,810 | 13,152,779 | 2.29 |
| 21 | 50,614,742 | 76,637 | 9,516,850 | 2.10 |
| 22 | 49,900,457 | 97,872 | 12,746,213 | 2.25 |
| $\sum$ or $\varnothing$ | **2,652,950,137** | **3,488,730** | **616,833,610** | **2.12** |

**Supplementary Table S2:** Summary of whole-genome, population-level data.

**Supplementary Figure S1**

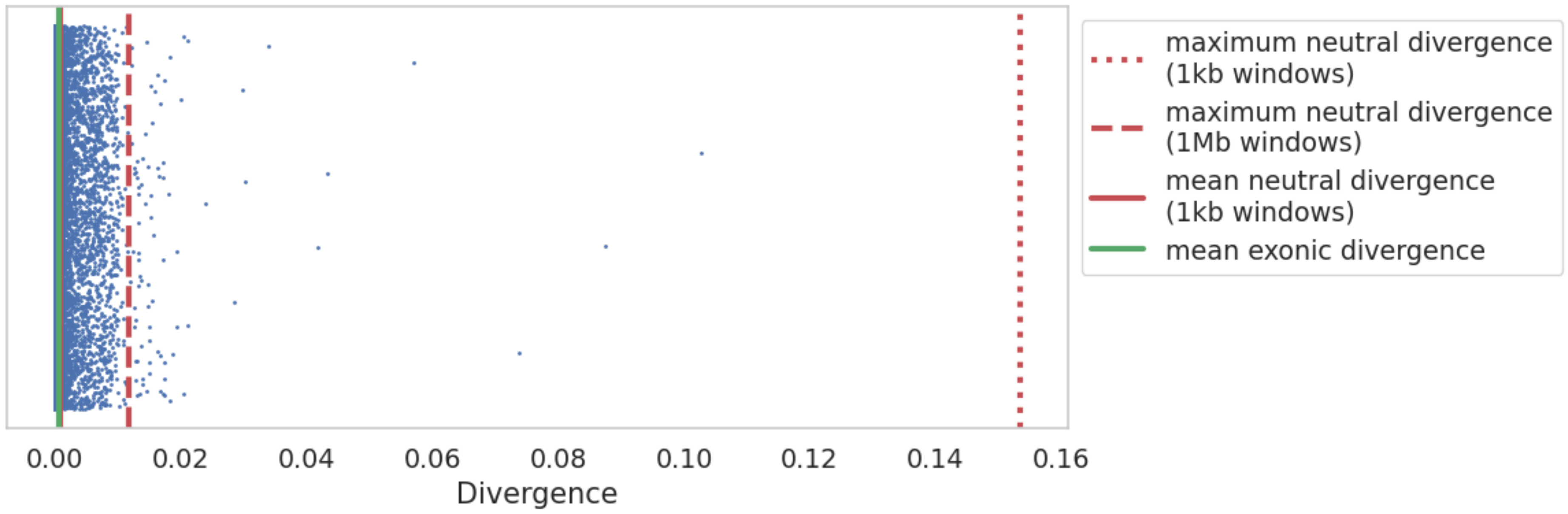


**Supplementary Figure S1:** Exonic divergence scatter plot with maximum neutral divergence values marked for windows of size 1Mb (red dashed line) and 1kb (red dotted line), as well as the mean neutral divergence for 1kb windows (red solid line), as calculated from Soni et al. (2025b). Each dot represents an autosomal exon, and the mean exonic divergence is plotted (green solid line).

**Supplementary Figure S2**

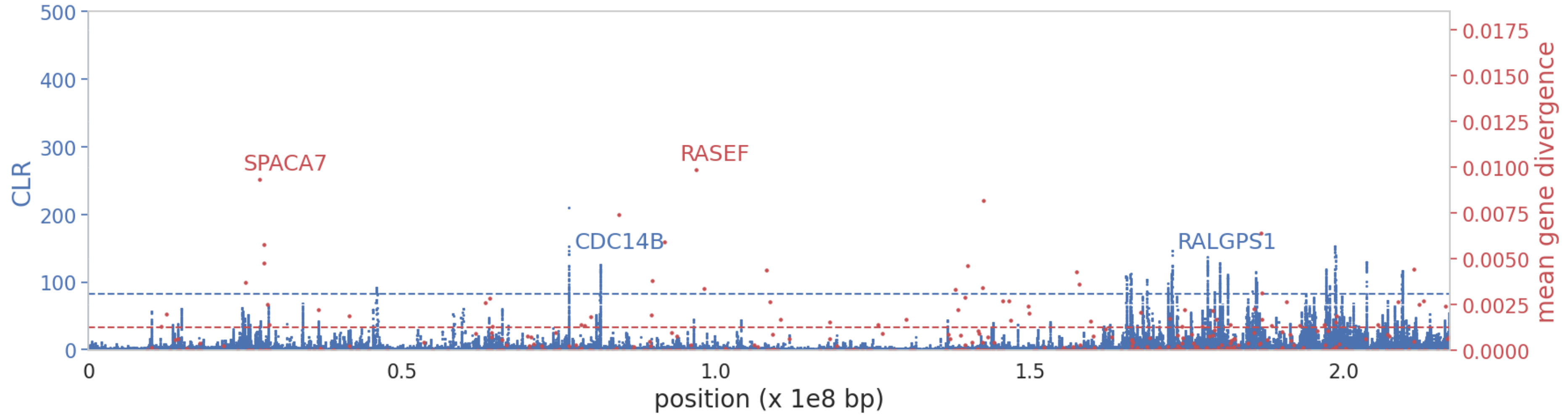


**Supplementary Figure S2:** SweepFinder2 selective sweep scan results (blue) and exonic divergence (red) for chromosome **1**. The x-axis represents the position along the chromosome, and the y-axis represents the CLR value at each SNP. Blue dashed line represents the null threshold for sweep detection; red dashed line represents 75$^{th}$ percentile neutral divergence. High CLR and highly divergent genes are marked by name.

**Supplementary Figure S3**

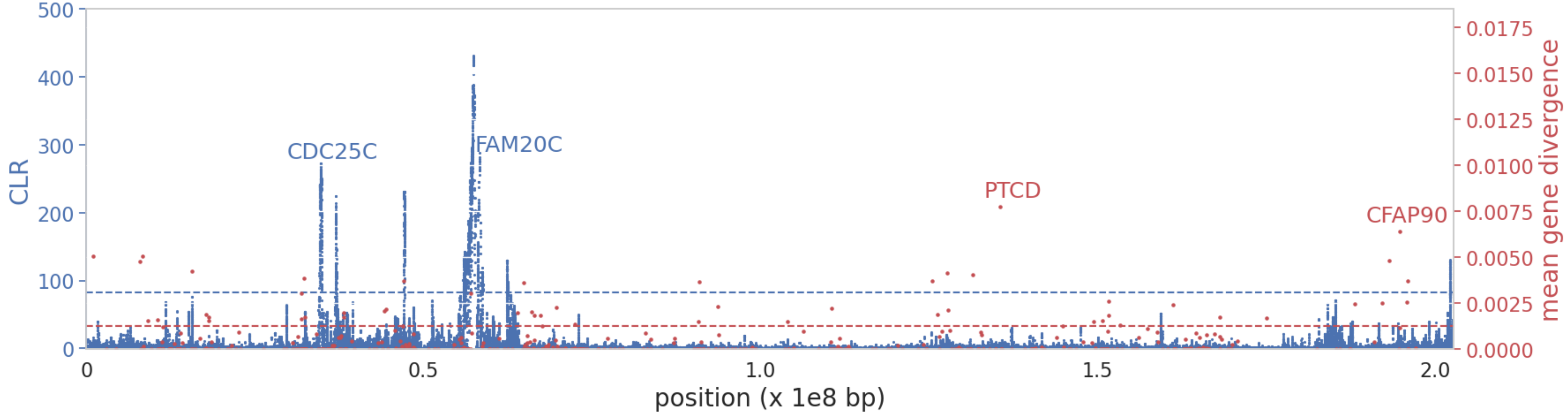


**Supplementary Figure S3:** SweepFinder2 selective sweep scan results (blue) and exonic divergence (red) for chromosome **2**. The x-axis represents the position along the chromosome, and the y-axis represents the CLR value at each SNP. Blue dashed line represents the null threshold for sweep detection; red dashed line represents 75th percentile neutral divergence. High CLR and highly divergent genes are marked by name.

**Supplementary Figure S4**

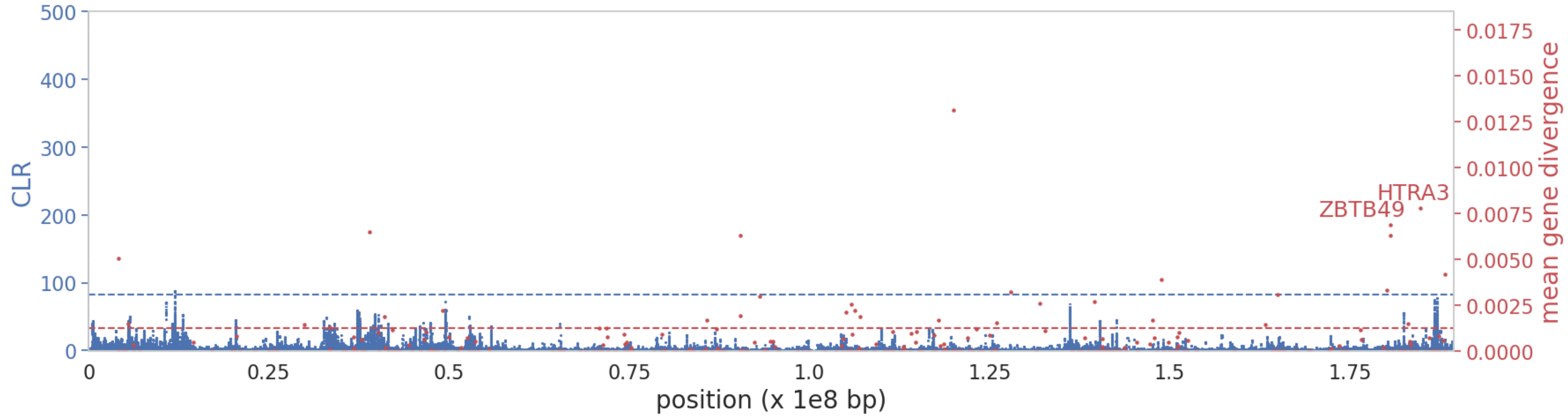


**Supplementary Figure S4:** SweepFinder2 selective sweep scan results (blue) and exonic divergence (red) for chromosome **3**. The x-axis represents the position along the chromosome, and the y-axis represents the CLR value at each SNP. Blue dashed line represents the null threshold for sweep detection; red dashed line represents 75th percentile neutral divergence. High CLR and highly divergent genes are marked by name.

**Supplementary Figure S5**

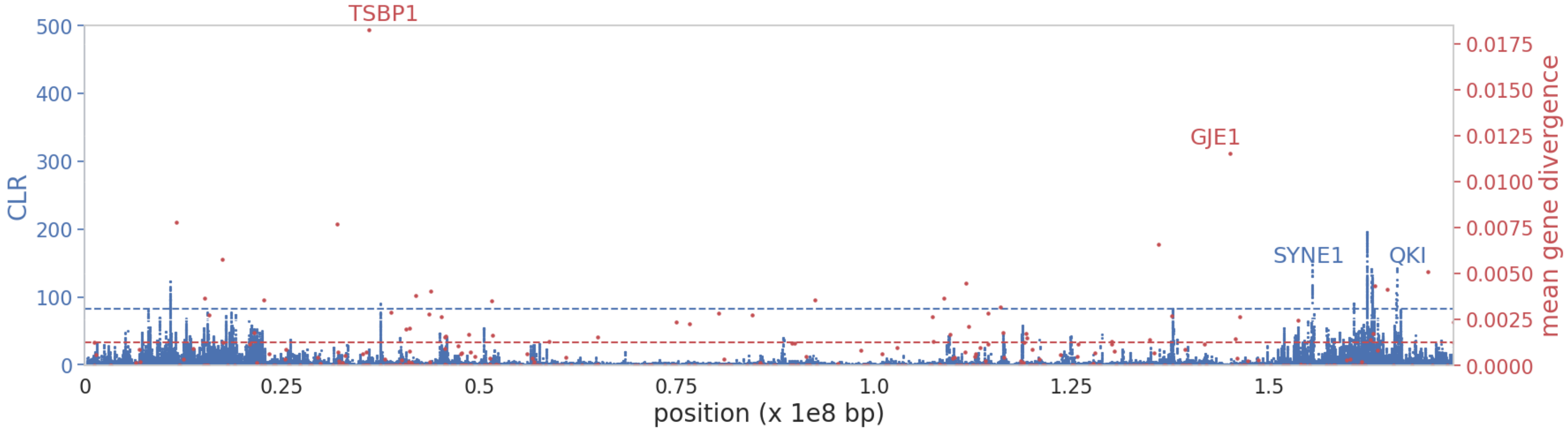


**Supplementary Figure S5:** SweepFinder2 selective sweep scan results (blue) and exonic divergence (red) for chromosome **4**. The x-axis represents the position along the chromosome, and the y-axis represents the CLR value at each SNP. Blue dashed line represents the null threshold for sweep detection; red dashed line represents 75th percentile neutral divergence. High CLR and highly divergent genes are marked by name.

**Supplementary Figure S6**

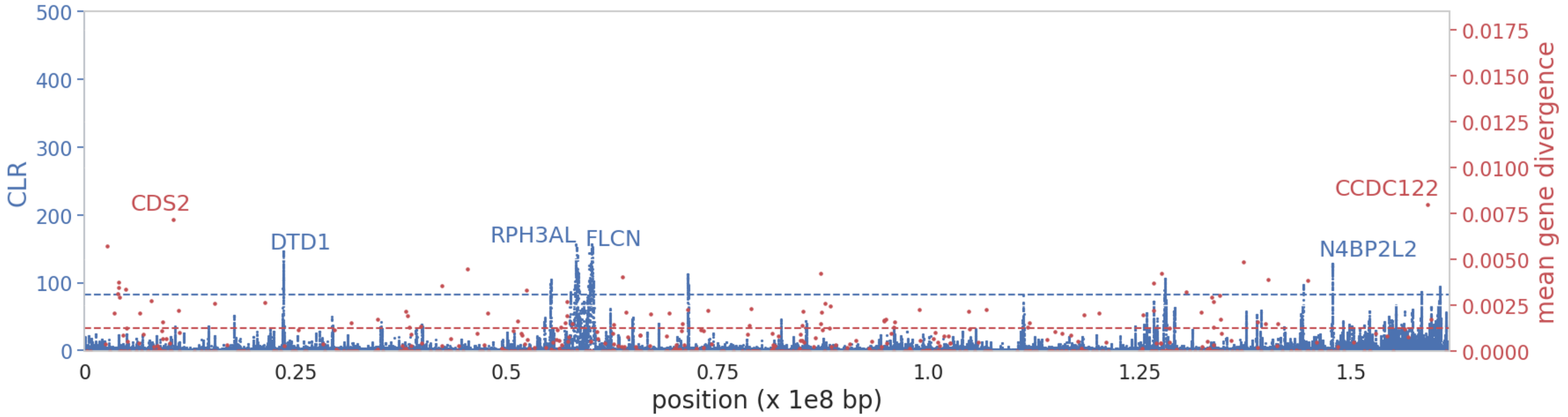


**Supplementary Figure S6:** SweepFinder2 selective sweep scan results (blue) and exonic divergence (red) for chromosome **5**. The x-axis represents the position along the chromosome, and the y-axis represents the CLR value at each SNP. Blue dashed line represents the null threshold for sweep detection; red dashed line represents 75$^{th}$ percentile neutral divergence. High CLR and highly divergent genes are marked by name.

**Supplementary Figure S7**

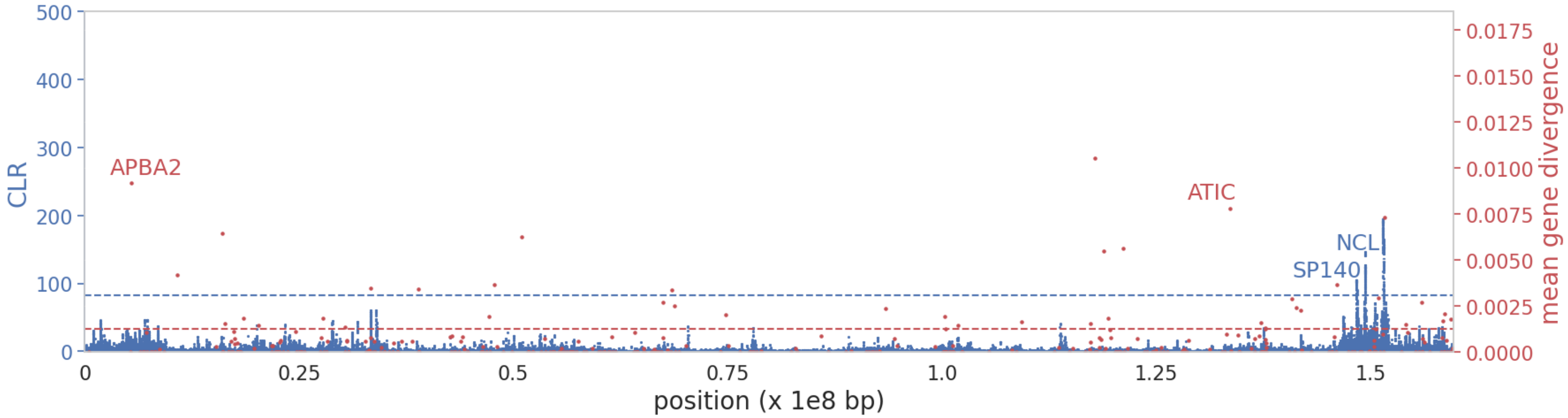


**Supplementary Figure S7:** SweepFinder2 selective sweep scan results (blue) and exonic divergence (red) for chromosome **6**. The x-axis represents the position along the chromosome, and the y-axis represents the CLR value at each SNP. Blue dashed line represents the null threshold for sweep detection; red dashed line represents 75$^{th}$ percentile neutral divergence. High CLR and highly divergent genes are marked by name.

**Supplementary Figure S8**

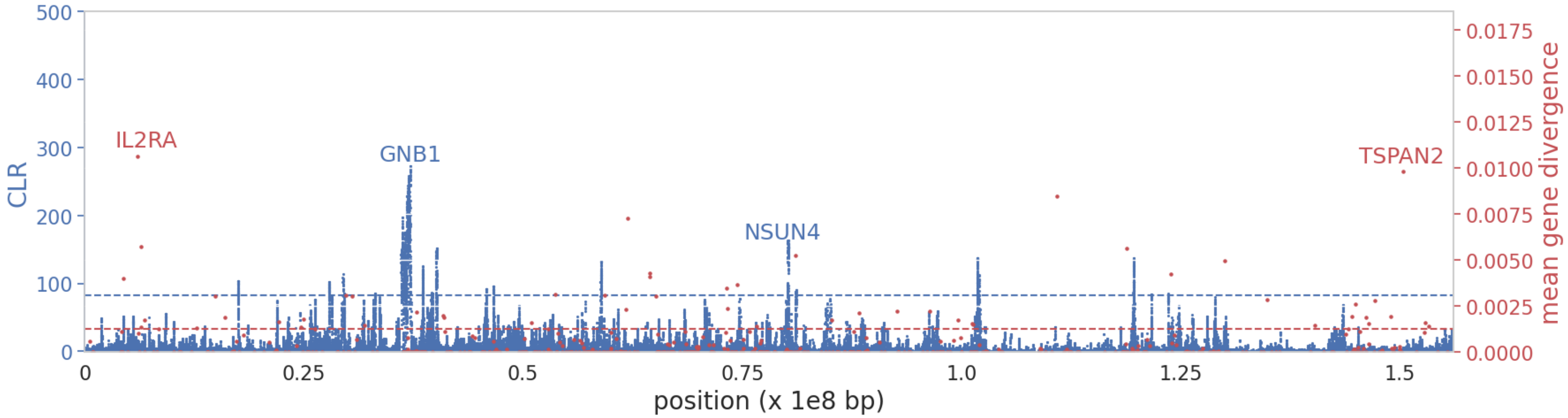


**Supplementary Figure S8:** SweepFinder2 selective sweep scan results (blue) and exonic divergence (red) for chromosome **7**. The x-axis represents the position along the chromosome, and the y-axis represents the CLR value at each SNP. Blue dashed line represents the null threshold for sweep detection; red dashed line represents 75$^{th}$ percentile neutral divergence. High CLR and highly divergent genes are marked by name.

**Supplementary Figure S9**

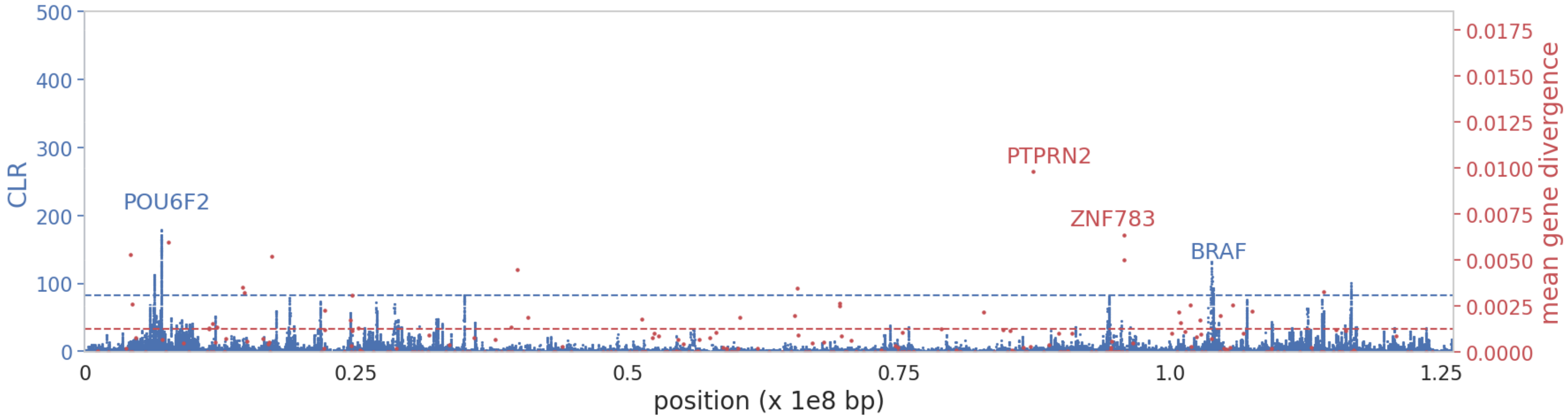


**Supplementary Figure S9:** SweepFinder2 selective sweep scan results (blue) and exonic divergence (red) for chromosome **8**. The x-axis represents the position along the chromosome, and the y-axis represents the CLR value at each SNP. Blue dashed line represents the null threshold for sweep detection; red dashed line represents 75$^{th}$ percentile neutral divergence. High CLR and highly divergent genes are marked by name.

**Supplementary Figure S10**

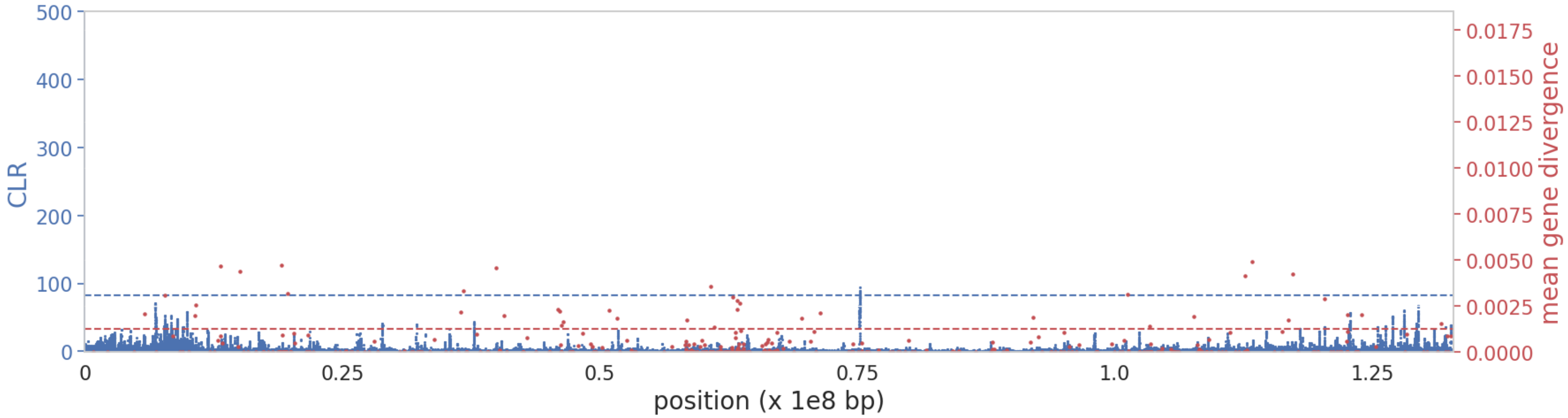


**Supplementary Figure S10:** SweepFinder2 selective sweep scan results (blue) and exonic divergence (red) for chromosome **9**. The x-axis represents the position along the chromosome, and the y-axis represents the CLR value at each SNP. Blue dashed line represents the null threshold for sweep detection; red dashed line represents 75th percentile neutral divergence. High CLR and highly divergent genes are marked by name.

**Supplementary Figure S11**

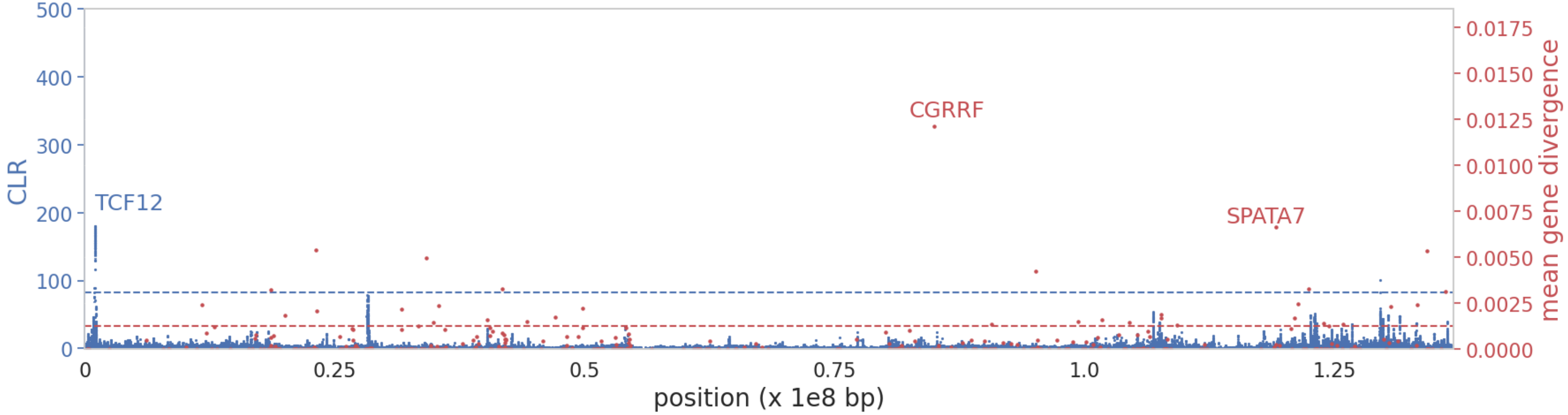


**Supplementary Figure S11:** SweepFinder2 selective sweep scan results (blue) and exonic divergence (red) for chromosome **10**. The x-axis represents the position along the chromosome, and the y-axis represents the CLR value at each SNP. Blue dashed line represents the null threshold for sweep detection; red dashed line represents 75th percentile neutral divergence. High CLR and highly divergent genes are marked by name.

**Supplementary Figure S12**

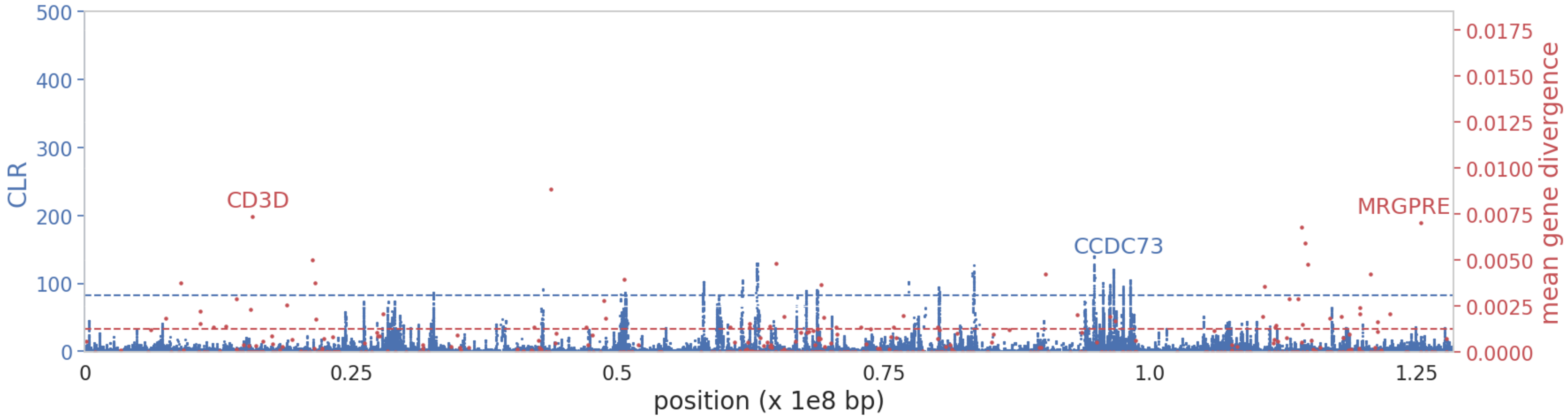


**Supplementary Figure S12:** SweepFinder2 selective sweep scan results (blue) and exonic divergence (red) for chromosome **11**. The x-axis represents the position along the chromosome, and the y-axis represents the CLR value at each SNP. Blue dashed line represents the null threshold for sweep detection; red dashed line represents 75$^{th}$ percentile neutral divergence. High CLR and highly divergent genes are marked by name.

**Supplementary Figure S13**

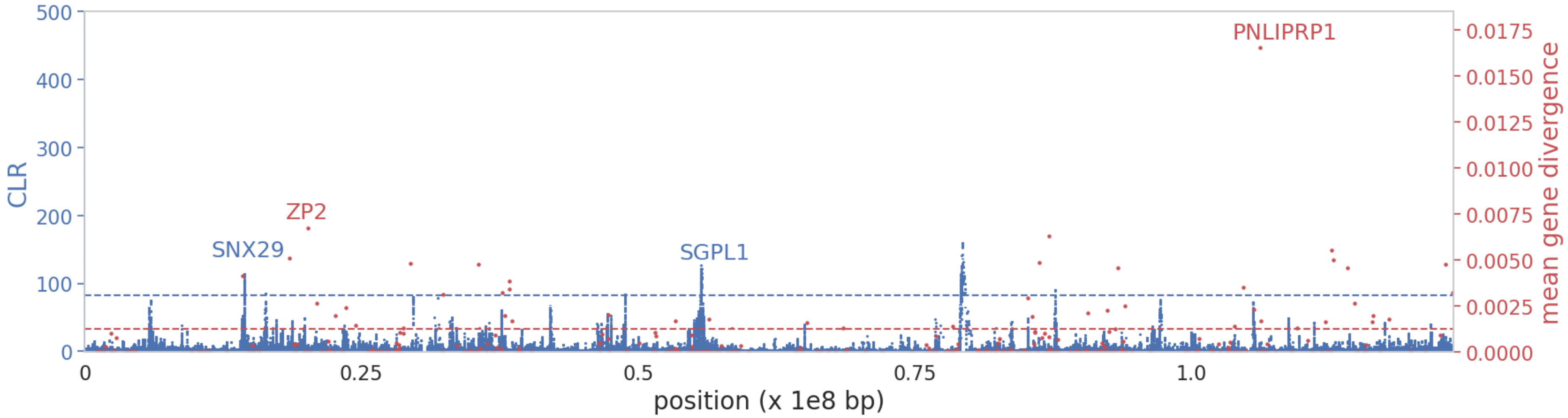


**Supplementary Figure S13:** SweepFinder2 selective sweep scan results (blue) and exonic divergence (red) for chromosome **12**. The x-axis represents the position along the chromosome, and the y-axis represents the CLR value at each SNP. Blue dashed line represents the null threshold for sweep detection; red dashed line represents 75th percentile neutral divergence. High CLR and highly divergent genes are marked by name.

**Supplementary Figure S14**

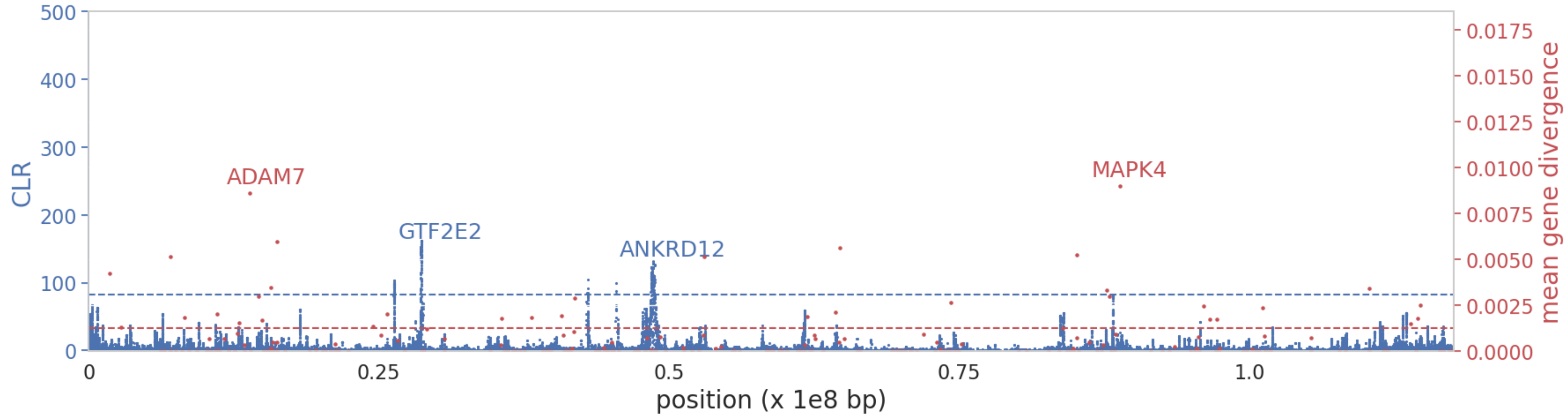


**Supplementary Figure S14:** SweepFinder2 selective sweep scan results (blue) and exonic divergence (red) for chromosome **13**. The x-axis represents the position along the chromosome, and the y-axis represents the CLR value at each SNP. Blue dashed line represents the null threshold for sweep detection; red dashed line represents 75th percentile neutral divergence. High CLR and highly divergent genes are marked by name.

**Supplementary Figure S15**

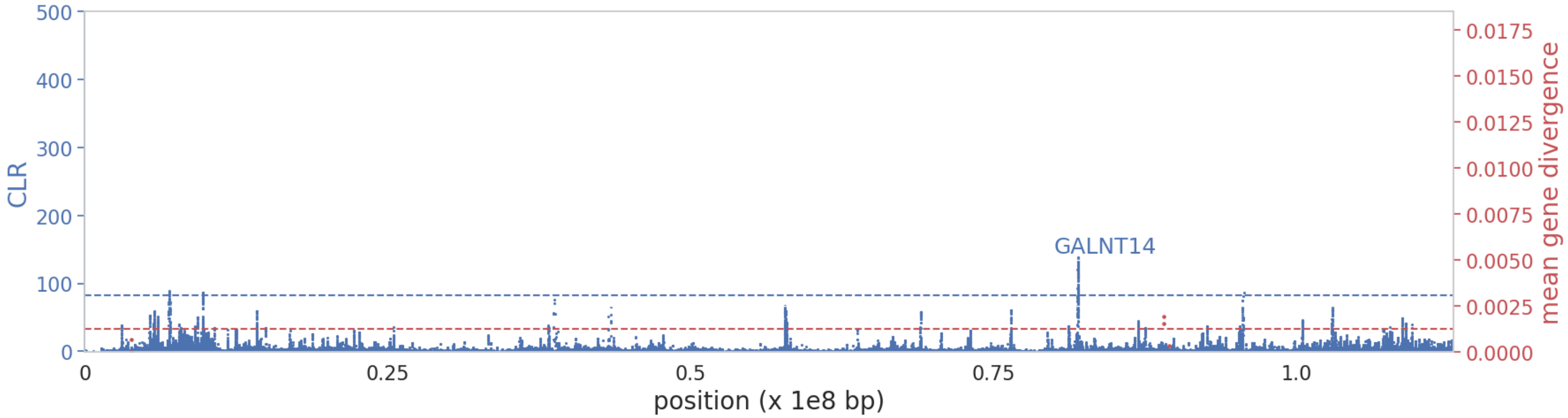


**Supplementary Figure S15:** SweepFinder2 selective sweep scan results (blue) and exonic divergence (red) for chromosome **14**. The x-axis represents the position along the chromosome, and the y-axis represents the CLR value at each SNP. Blue dashed line represents the null threshold for sweep detection; red dashed line represents 75th percentile neutral divergence. High CLR and highly divergent genes are marked by name.

**Supplementary Figure S16**

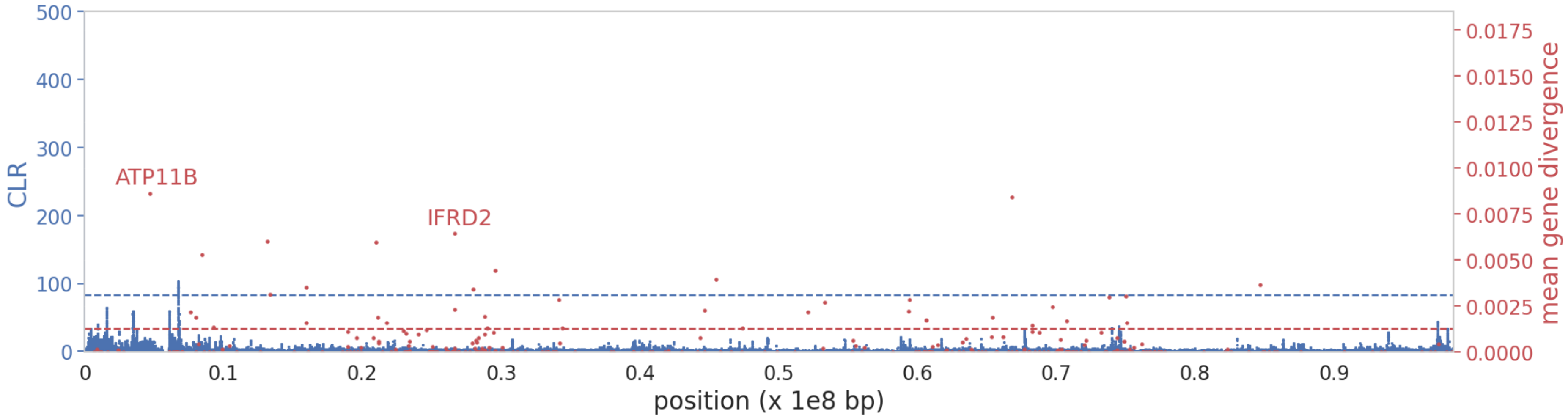


**Supplementary Figure S16:** SweepFinder2 selective sweep scan results (blue) and exonic divergence (red) for chromosome **15**. The x-axis represents the position along the chromosome, and the y-axis represents the CLR value at each SNP. Blue dashed line represents the null threshold for sweep detection; red dashed line represents 75$^{th}$ percentile neutral divergence. High CLR and highly divergent genes are marked by name.

**Supplementary Figure S17**

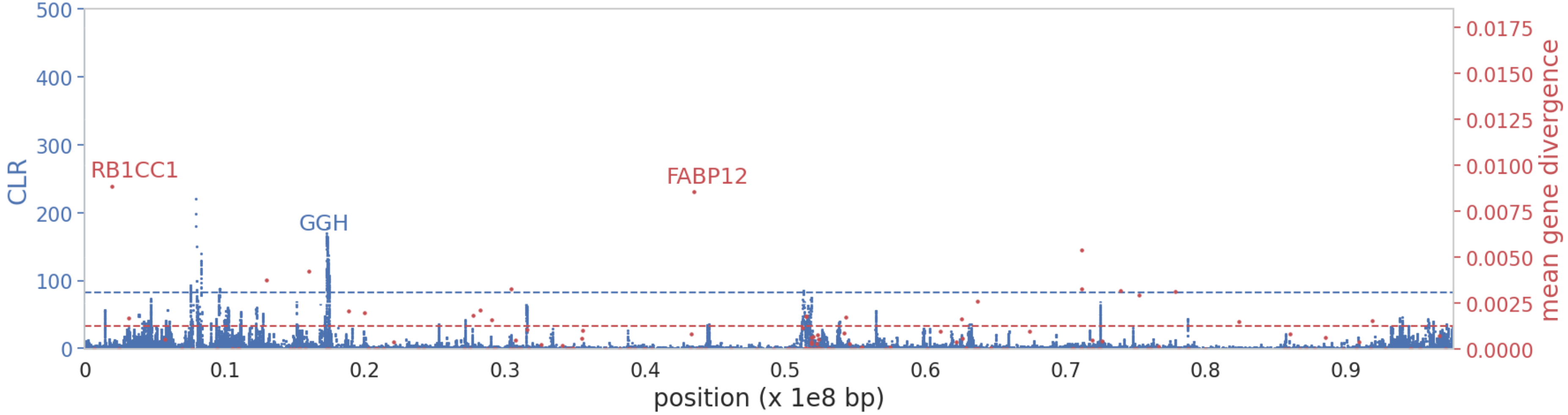


**Supplementary Figure S17:** SweepFinder2 selective sweep scan results (blue) and exonic divergence (red) for chromosome **16**. The x-axis represents the position along the chromosome, and the y-axis represents the CLR value at each SNP. Blue dashed line represents the null threshold for sweep detection; red dashed line represents 75th percentile neutral divergence. High CLR and highly divergent genes are marked by name.

**Supplementary Figure S18**

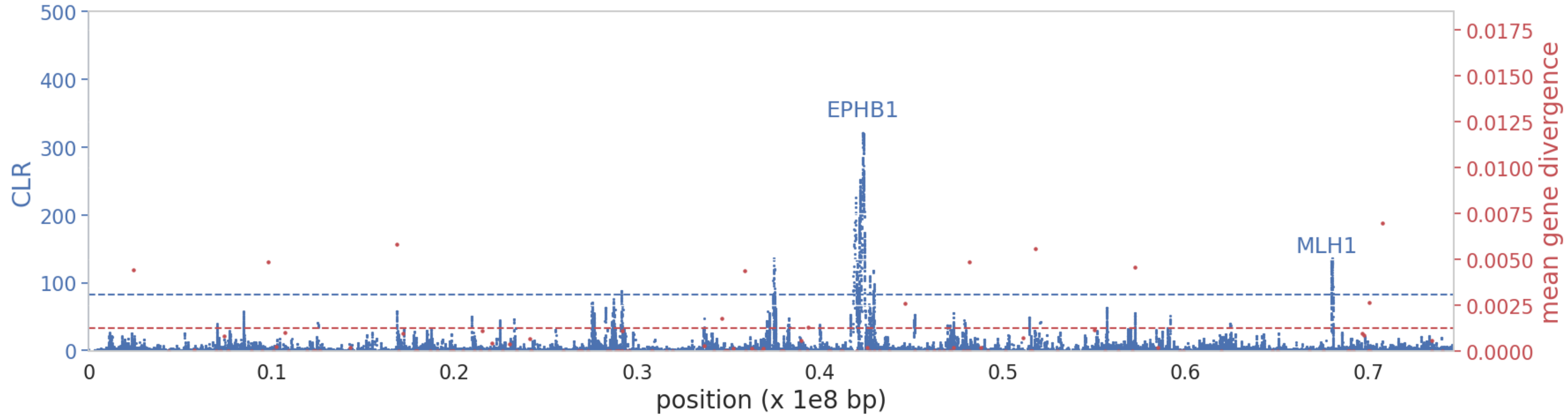


**Supplementary Figure S18:** SweepFinder2 selective sweep scan results (blue) and exonic divergence (red) for chromosome **17**. The x-axis represents the position along the chromosome, and the y-axis represents the CLR value at each SNP. Blue dashed line represents the null threshold for sweep detection; red dashed line represents 75th percentile neutral divergence. High CLR and highly divergent genes are marked by name.

**Supplementary Figure S19**

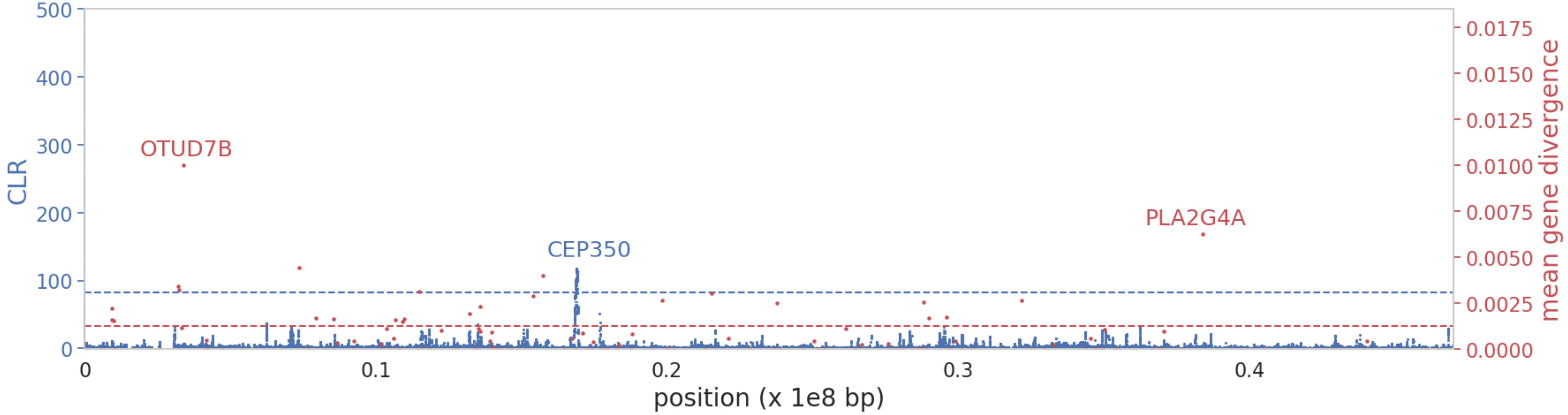


**Supplementary Figure S19:** SweepFinder2 selective sweep scan results (blue) and exonic divergence (red) for chromosome **18**. The x-axis represents the position along the chromosome, and the y-axis represents the CLR value at each SNP. Blue dashed line represents the null threshold for sweep detection; red dashed line represents 75th percentile neutral divergence. High CLR and highly divergent genes are marked by name.

**Supplementary Figure S20**

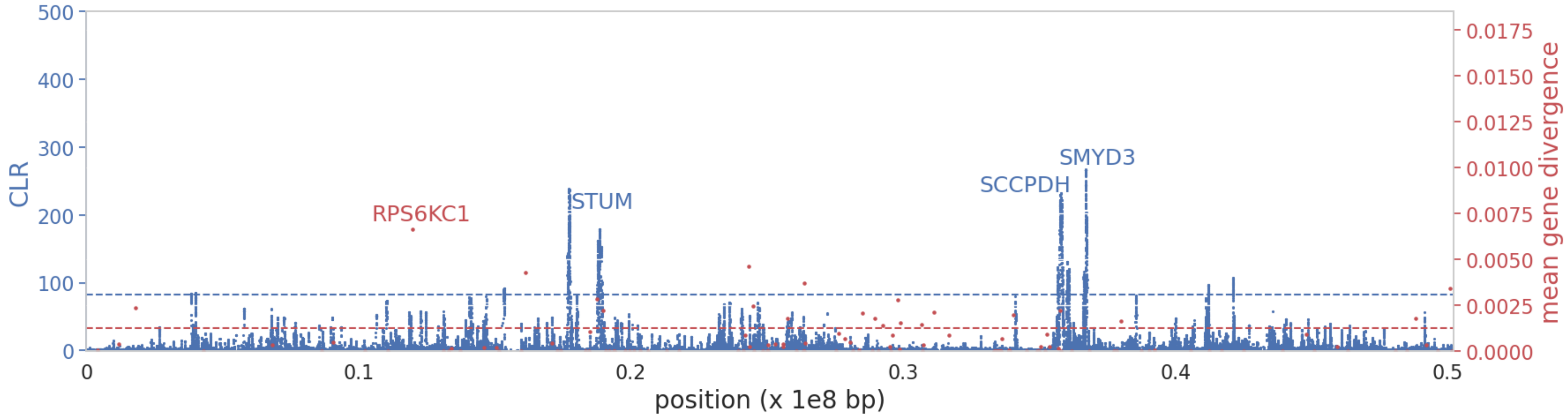


**Supplementary Figure S20:** SweepFinder2 selective sweep scan results (blue) and exonic divergence (red) for chromosome **19**. The x-axis represents the position along the chromosome, and the y-axis represents the CLR value at each SNP. Blue dashed line represents the null threshold for sweep detection; red dashed line represents 75$^{th}$ percentile neutral divergence. High CLR and highly divergent genes are marked by name.

**Supplementary Figure S21**

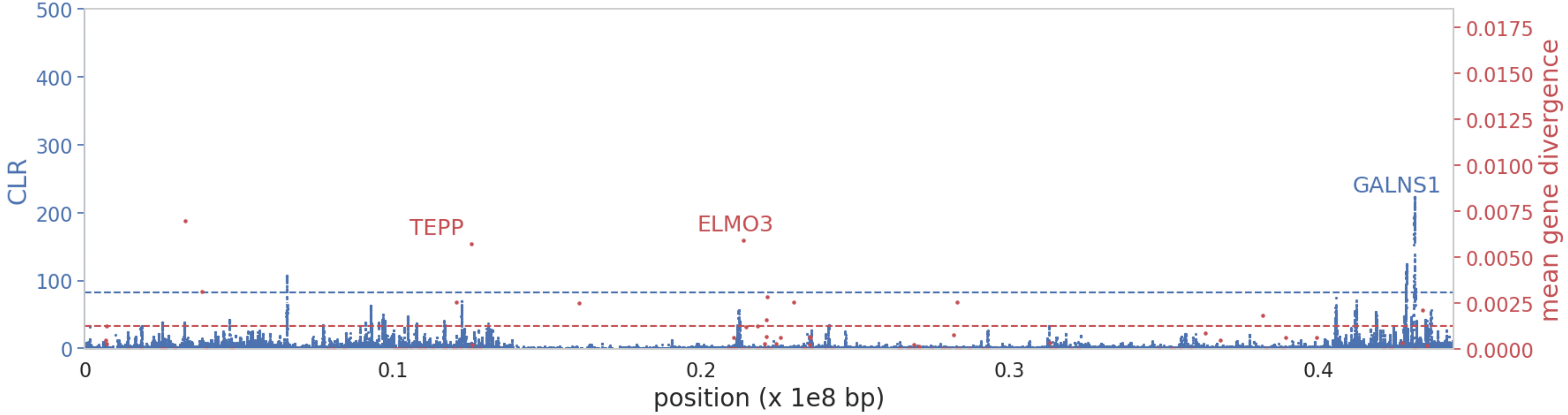


**Supplementary Figure S21:** SweepFinder2 selective sweep scan results (blue) and exonic divergence (red) for chromosome **20**. The x-axis represents the position along the chromosome, and the y-axis represents the CLR value at each SNP. Blue dashed line represents the null threshold for sweep detection; red dashed line represents 75$^{th}$ percentile neutral divergence. High CLR and highly divergent genes are marked by name.

**Supplementary Figure S22**

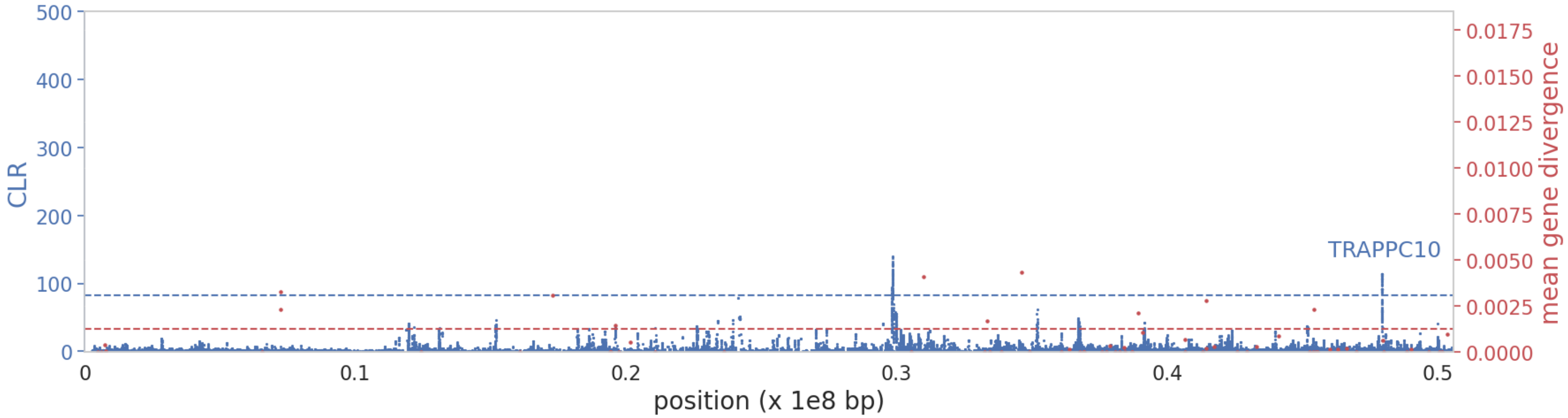


**Supplementary Figure S22:** SweepFinder2 selective sweep scan results (blue) and exonic divergence (red) for chromosome **21**. The x-axis represents the position along the chromosome, and the y-axis represents the CLR value at each SNP. Blue dashed line represents the null threshold for sweep detection; red dashed line represents 75$^{th}$ percentile neutral divergence. High CLR and highly divergent genes are marked by name.

**Supplementary Figure S23**

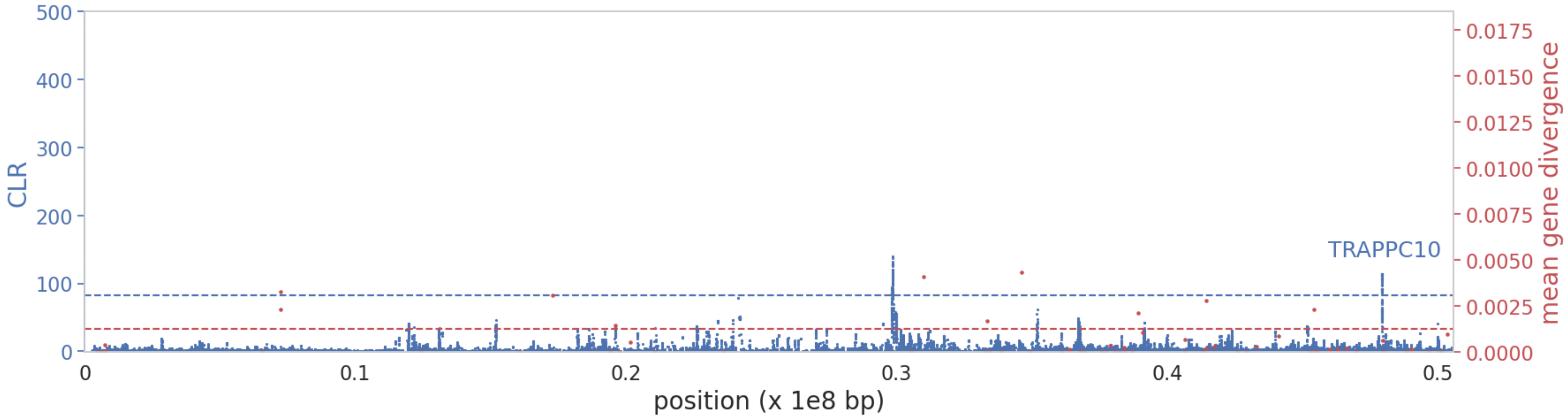


**Supplementary Figure S23:** SweepFinder2 selective sweep scan results (blue) and exonic divergence (red) for chromosome **22**. The x-axis represents the position along the chromosome, and the y-axis represents the CLR value at each SNP. Blue dashed line represents the null threshold for sweep detection; red dashed line represents 75th percentile neutral divergence. High CLR and highly divergent genes are marked by name.

**Supplementary Figure S24**

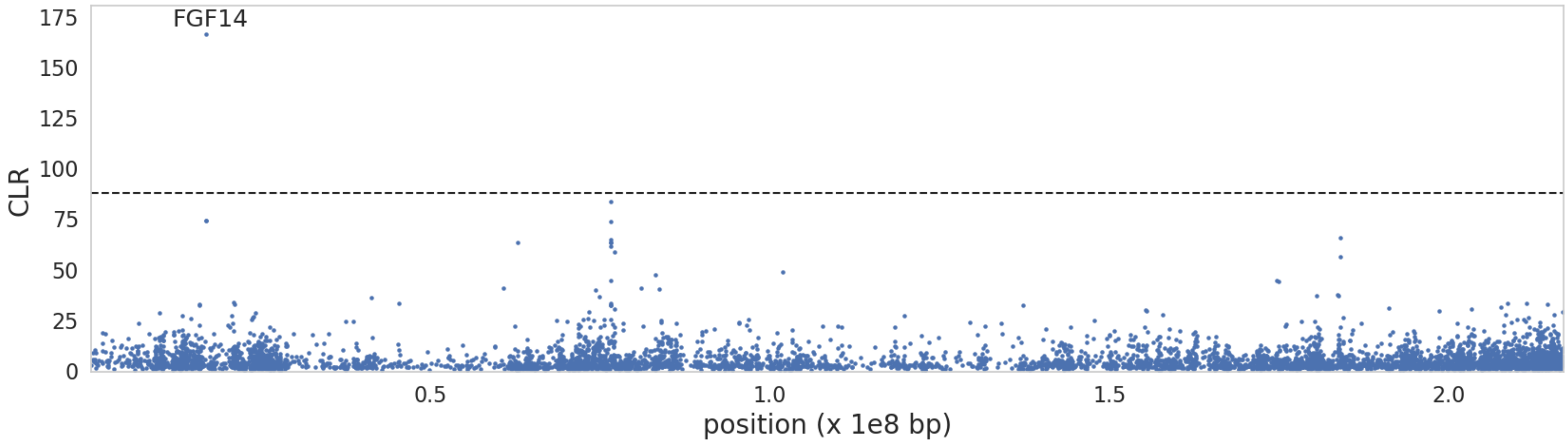


**Supplementary Figure S24:** $B_{0MAF}$ balancing selection scan results for chromosome **1**. The x-axis represents the position along the chromosome, and the y-axis represents the CLR value at each 1kb window (with a 50bp step size). Black dashed line represents the null threshold for balancing selection inference. Candidate genes are labelled on plot.

**Supplementary Figure S25**

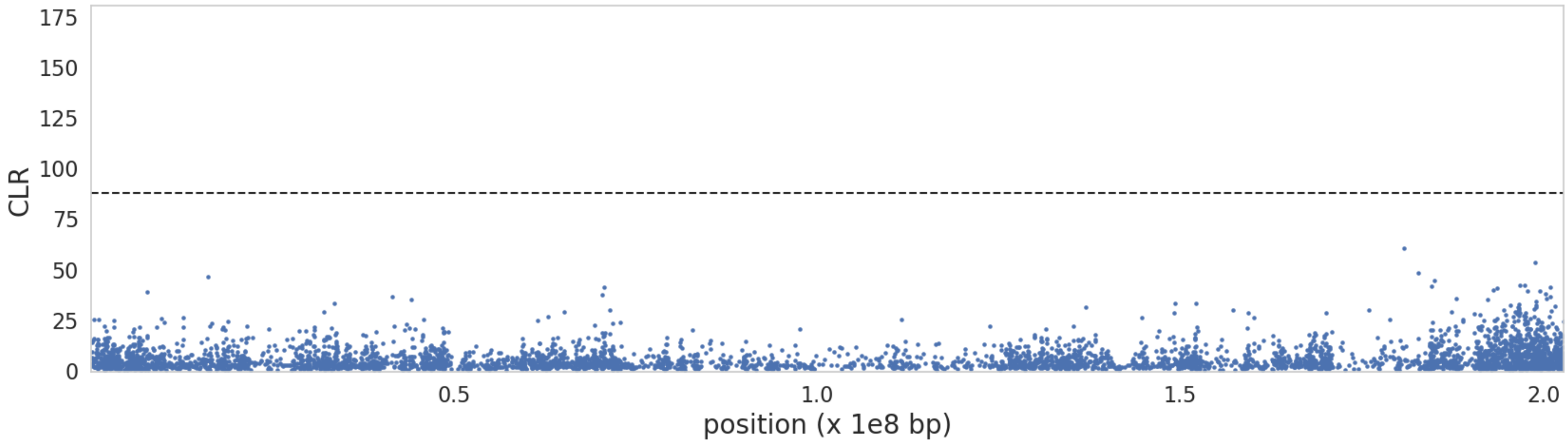


**Supplementary Figure S25:** $B_{0MAF}$ balancing selection scan results for chromosome **2**. The x-axis represents the position along the chromosome, and the y-axis represents the CLR value at each 1kb window (with a 50bp step size). Black dashed line represents the null threshold for balancing selection inference. Candidate genes are labelled on plot.

**Supplementary Figure S26**

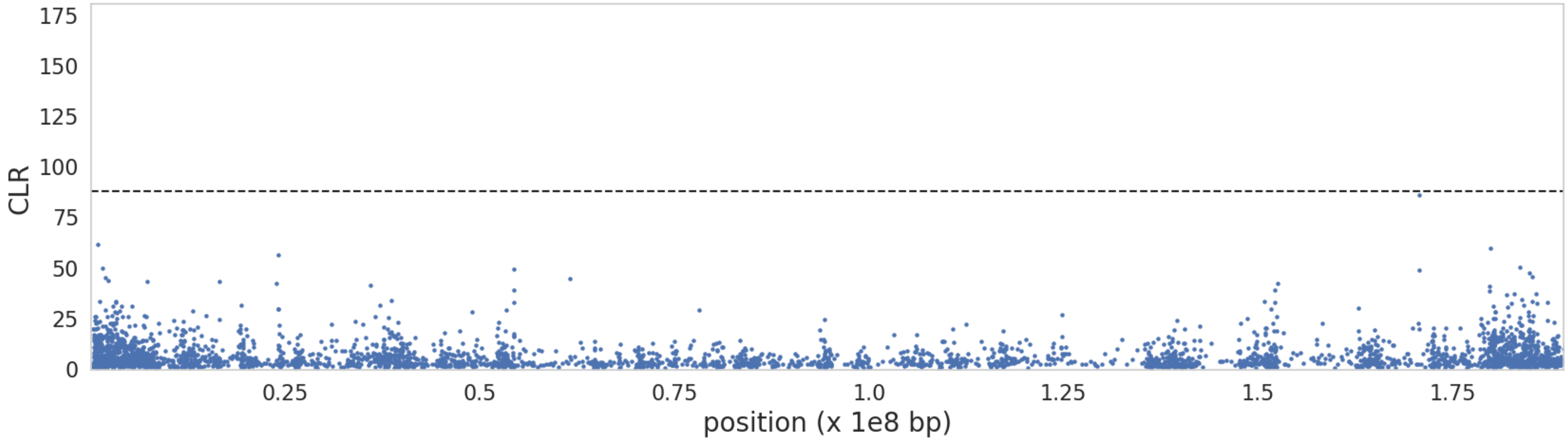


**Supplementary Figure S26:** $B_{0MAF}$ balancing selection scan results for chromosome **3**. The x-axis represents the position along the chromosome, and the y-axis represents the CLR value at each 1kb window (with a 50bp step size). Black dashed line represents the null threshold for balancing selection inference. Candidate genes are labelled on plot.

**Supplementary Figure S27**

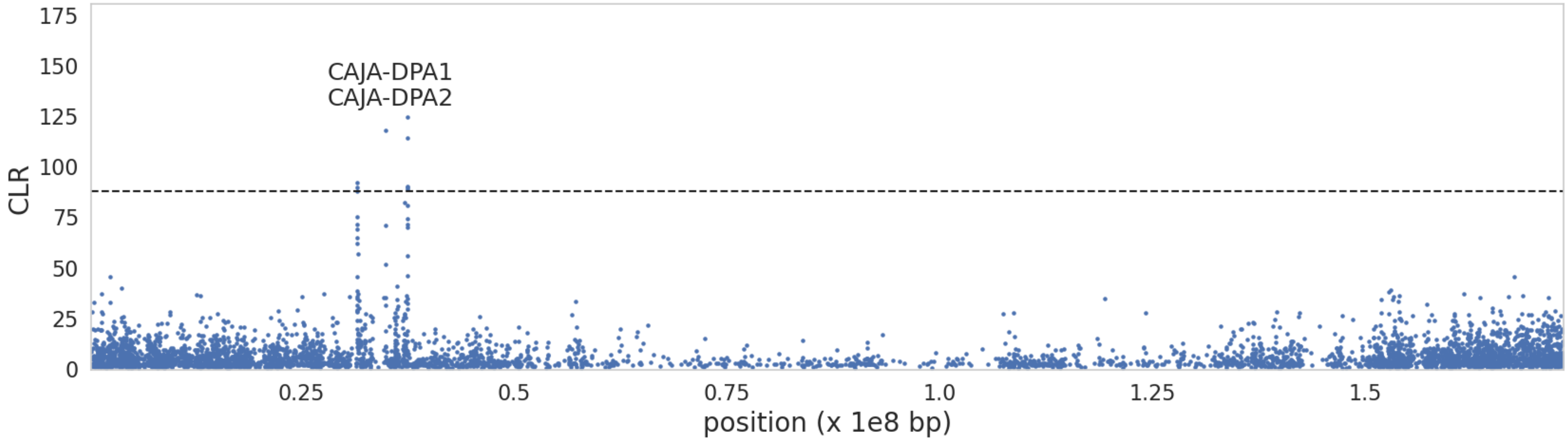


**Supplementary Figure S27:** $B_{0MAF}$ balancing selection scan results for chromosome **4**. The x-axis represents the position along the chromosome, and the y-axis represents the CLR value at each 1kb window (with a 50bp step size). Black dashed line represents the null threshold for balancing selection inference. Candidate genes are labelled on plot.

**Supplementary Figure S28**

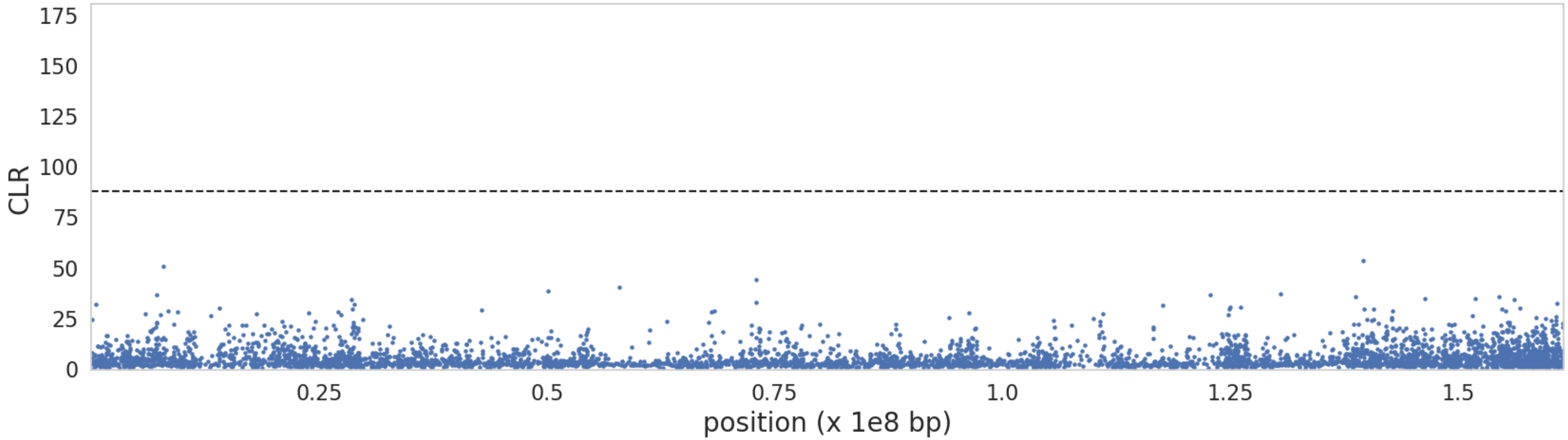


**Supplementary Figure S28:** $B_{0MAF}$ balancing selection scan results for chromosome **5**. The x-axis represents the position along the chromosome, and the y-axis represents the CLR value at each 1kb window (with a 50bp step size). Black dashed line represents the null threshold for balancing selection inference. Candidate genes are labelled on plot.

**Supplementary Figure S29**

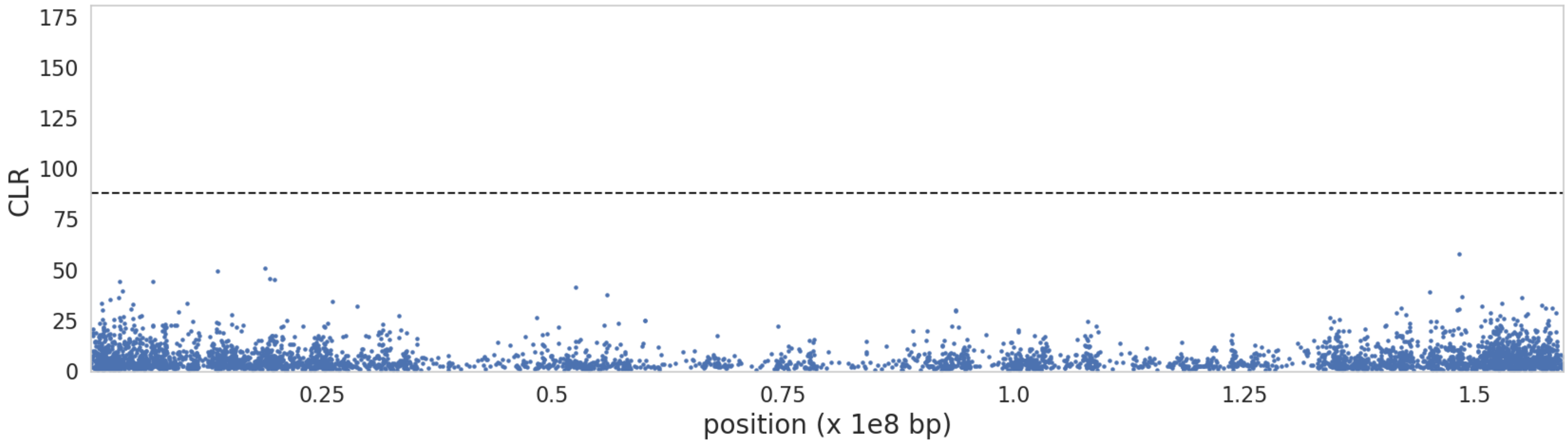


**Supplementary Figure S29:** $B_{0MAF}$ balancing selection scan results for chromosome **6**. The x-axis represents the position along the chromosome, and the y-axis represents the CLR value at each 1kb window (with a 50bp step size). Black dashed line represents the null threshold for balancing selection inference. Candidate genes are labelled on plot.

**Supplementary Figure S30**

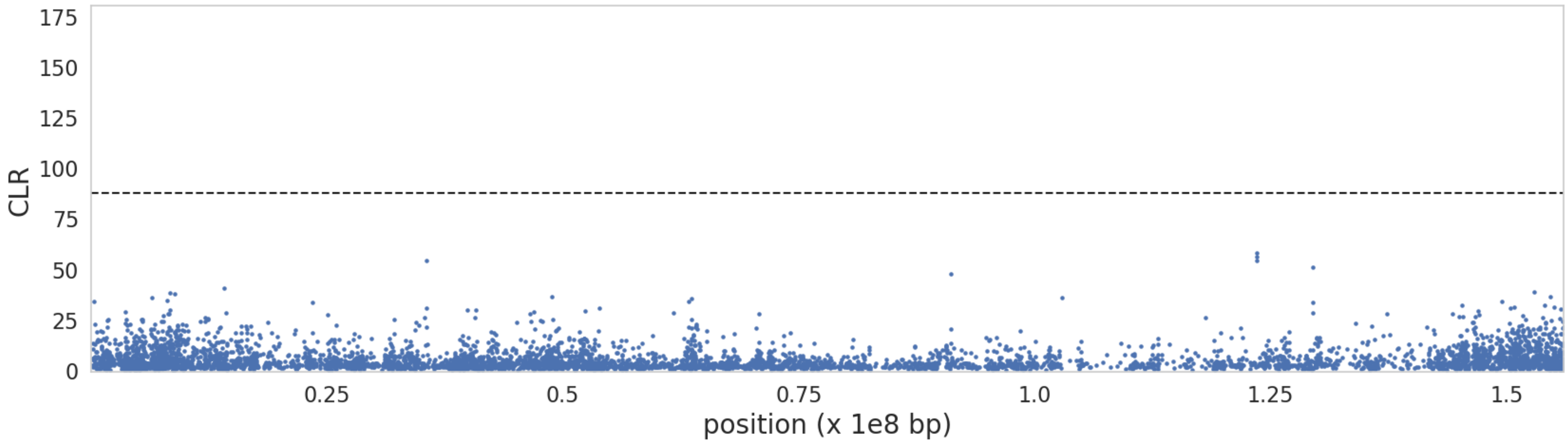


**Supplementary Figure S30:** $B_{0MAF}$ balancing selection scan results for chromosome **7**. The x-axis represents the position along the chromosome, and the y-axis represents the CLR value at each 1kb window (with a 50bp step size). Black dashed line represents the null threshold for balancing selection inference. Candidate genes are labelled on plot.

**Supplementary Figure S31**

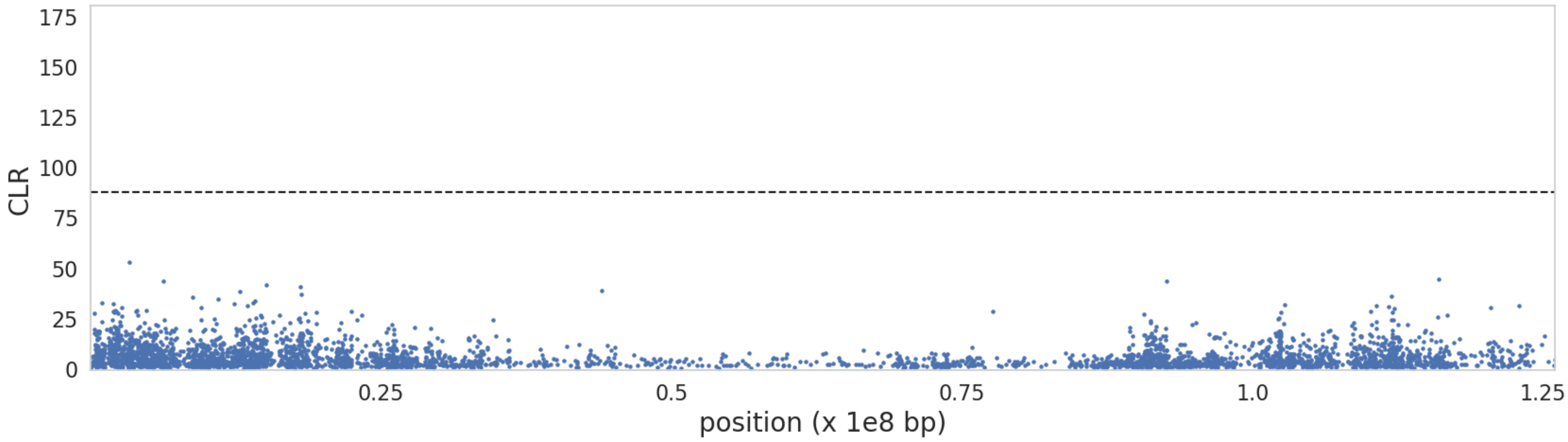


**Supplementary Figure S31:** $B_{0MAF}$ balancing selection scan results for chromosome **8**. The x-axis represents the position along the chromosome, and the y-axis represents the CLR value at each 1kb window (with a 50bp step size). Black dashed line represents the null threshold for balancing selection inference. Candidate genes are labelled on plot.

**Supplementary Figure S32**

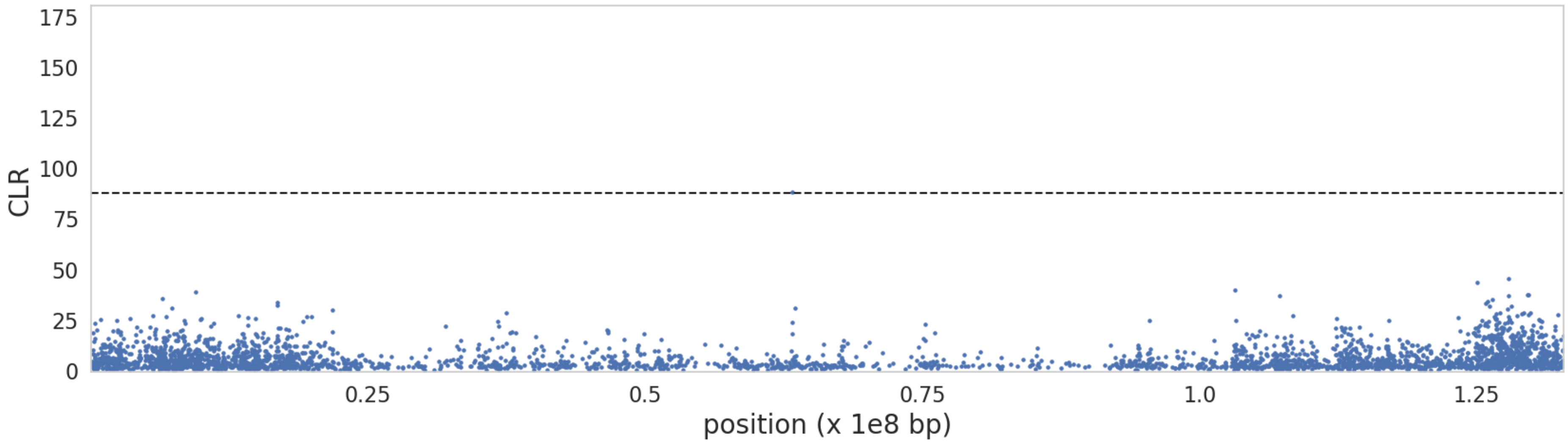


**Supplementary Figure S32:** $B_{0MAF}$ balancing selection scan results for chromosome **9**. The x-axis represents the position along the chromosome, and the y-axis represents the CLR value at each 1kb window (with a 50bp step size). Black dashed line represents the null threshold for balancing selection inference. Candidate genes are labelled on plot.

**Supplementary Figure S33**

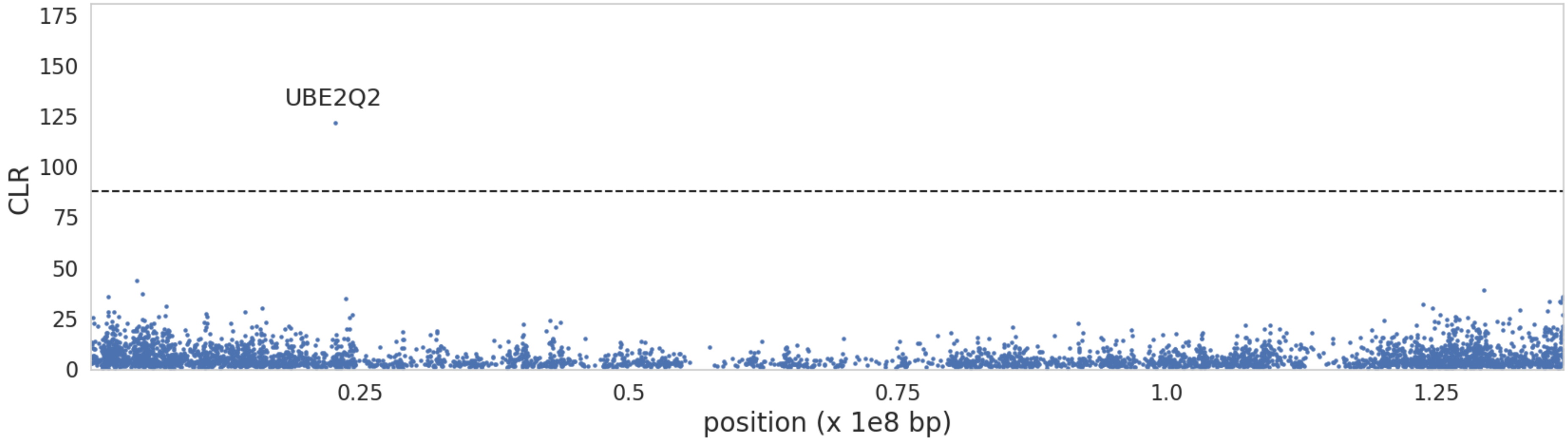


**Supplementary Figure S33:** $B_{0MAF}$ balancing selection scan results for chromosome **10**. The x-axis represents the position along the chromosome, and the y-axis represents the CLR value at each 1kb window (with a 50bp step size). Black dashed line represents the null threshold for balancing selection inference. Candidate genes are labelled on plot.

**Supplementary Figure S34**

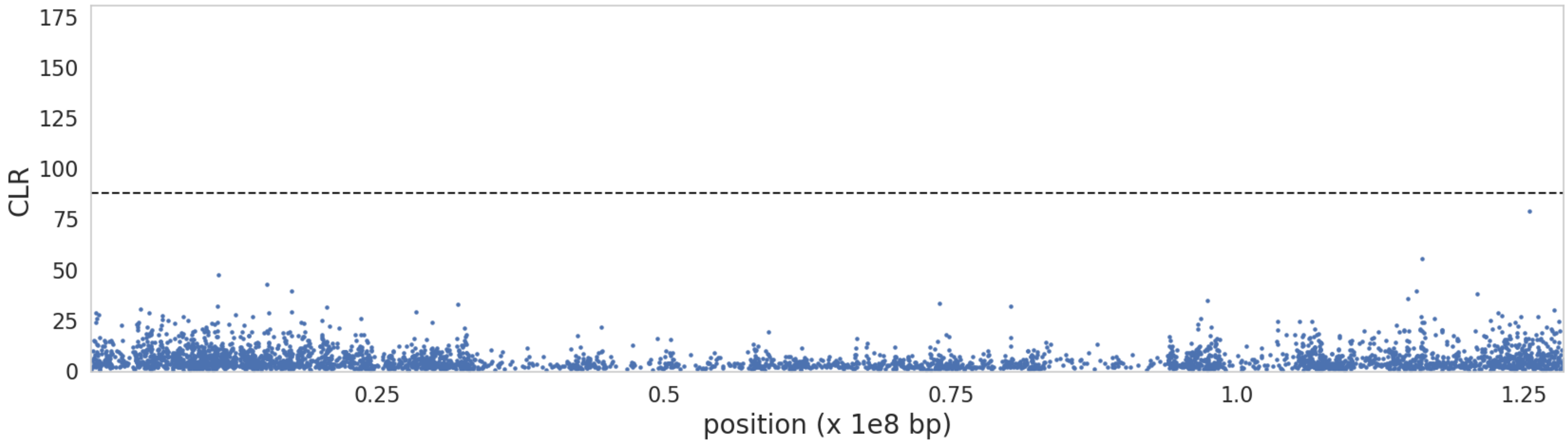


**Supplementary Figure S34:** $B_{0MAF}$ balancing selection scan results for chromosome **11**. The x-axis represents the position along the chromosome, and the y-axis represents the CLR value at each 1kb window (with a 50bp step size). Black dashed line represents the null threshold for balancing selection inference. Candidate genes are labelled on plot.

**Supplementary Figure S35**

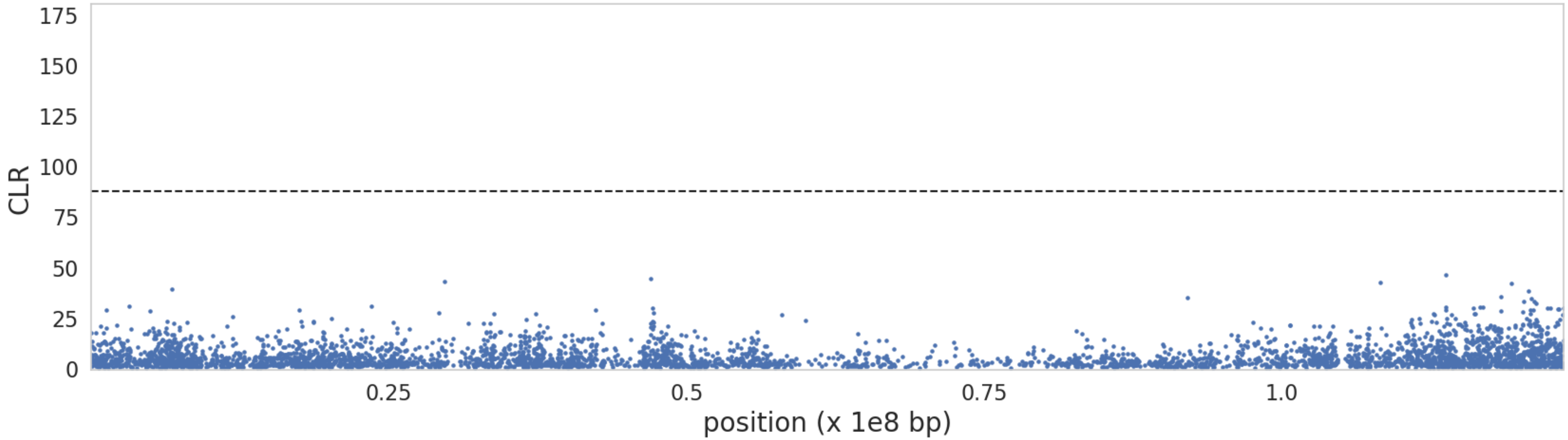


**Supplementary Figure S35:** $B_{0MAF}$ balancing selection scan results for chromosome **12**. The x-axis represents the position along the chromosome, and the y-axis represents the CLR value at each 1kb window (with a 50bp step size). Black dashed line represents the null threshold for balancing selection inference. Candidate genes are labelled on plot.

**Supplementary Figure S36**

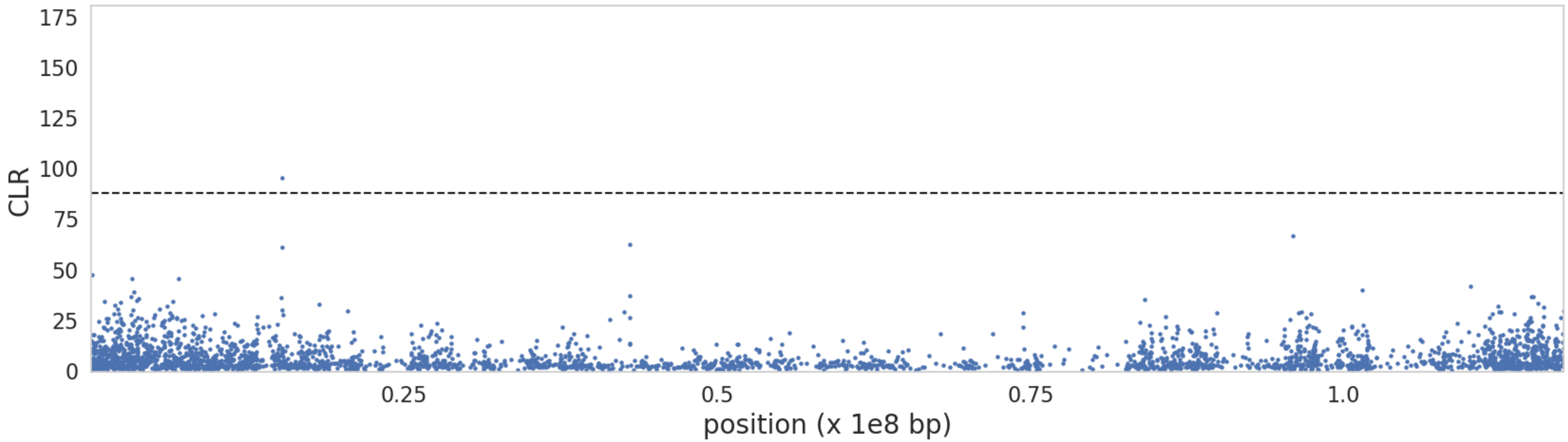


**Supplementary Figure S36:** $B_{0MAF}$ balancing selection scan results for chromosome **13**. The x-axis represents the position along the chromosome, and the y-axis represents the CLR value at each 1kb window (with a 50bp step size). Black dashed line represents the null threshold for balancing selection inference. Candidate genes are labelled on plot.

**Supplementary Figure S37**

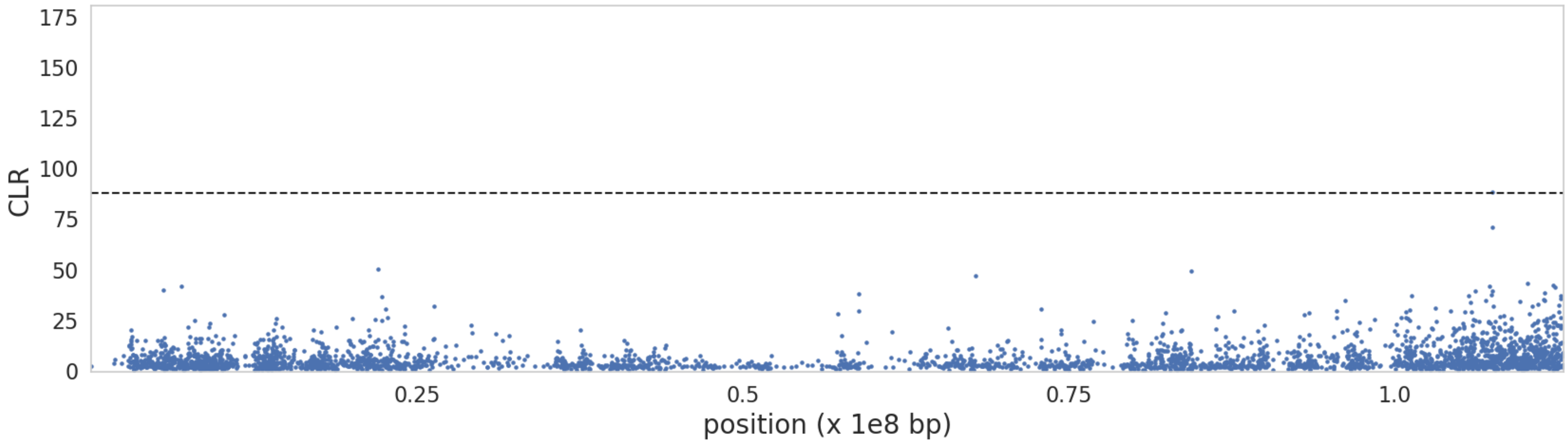


**Supplementary Figure S37:** $B_{0MAF}$ balancing selection scan results for chromosome **14**. The x-axis represents the position along the chromosome, and the y-axis represents the CLR value at each 1kb window (with a 50bp step size). Black dashed line represents the null threshold for balancing selection inference. Candidate genes are labelled on plot.

**Supplementary Figure S38**

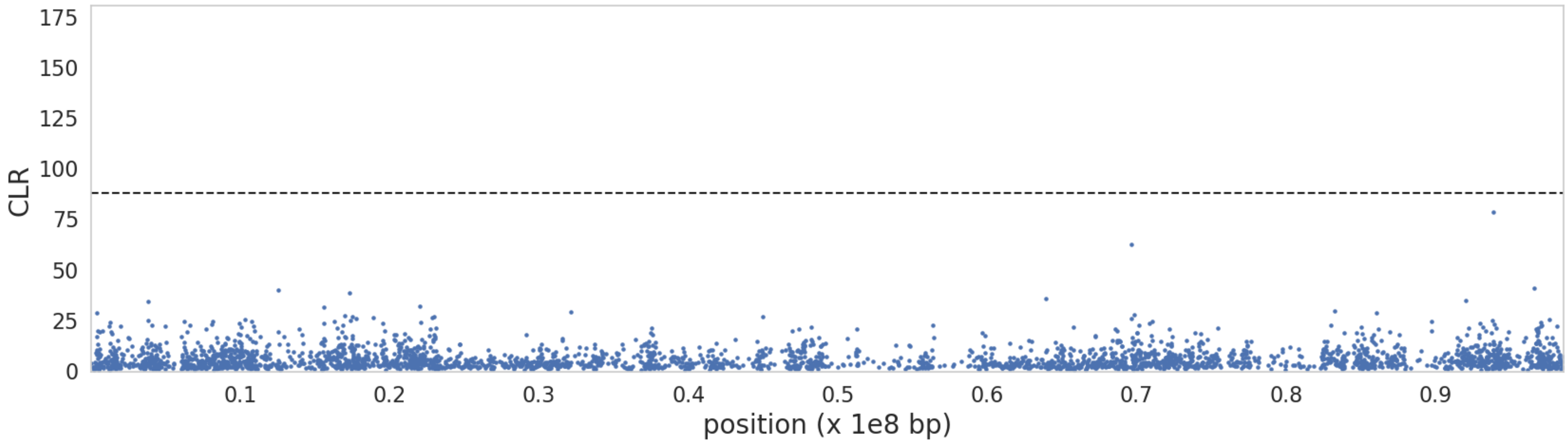


**Supplementary Figure S38:** $B_{0MAF}$ balancing selection scan results for chromosome **15**. The x-axis represents the position along the chromosome, and the y-axis represents the CLR value at each 1kb window (with a 50bp step size). Black dashed line represents the null threshold for balancing selection inference. Candidate genes are labelled on plot.

**Supplementary Figure S39**

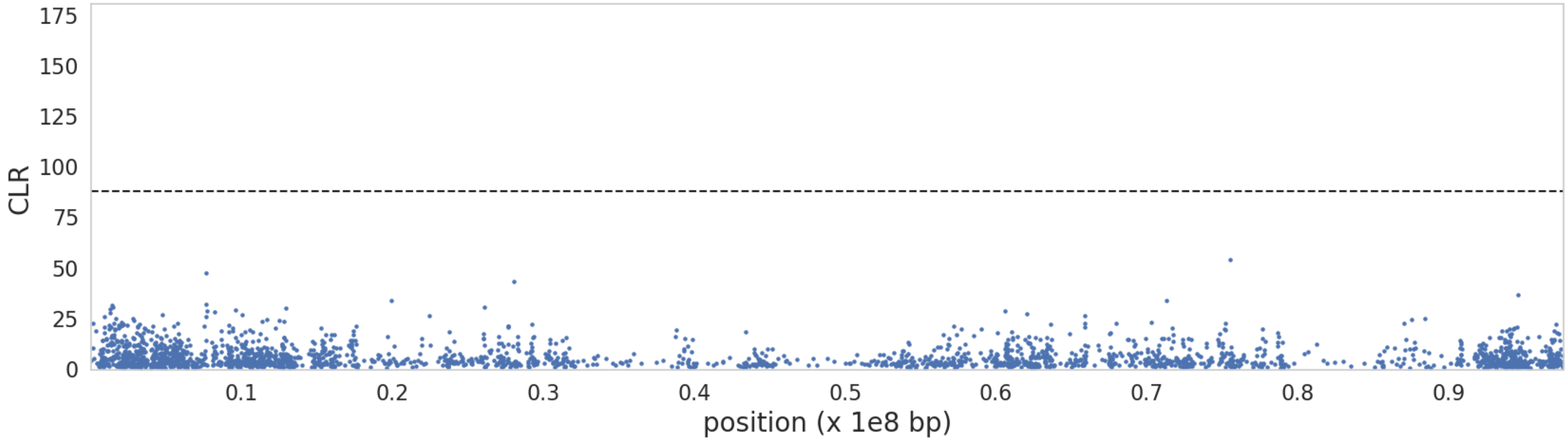


**Supplementary Figure S39:** $B_{0MAF}$ balancing selection scan results for chromosome **16**. The x-axis represents the position along the chromosome, and the y-axis represents the CLR value at each 1kb window (with a 50bp step size). Black dashed line represents the null threshold for balancing selection inference. Candidate genes are labelled on plot.

**Supplementary Figure S40**

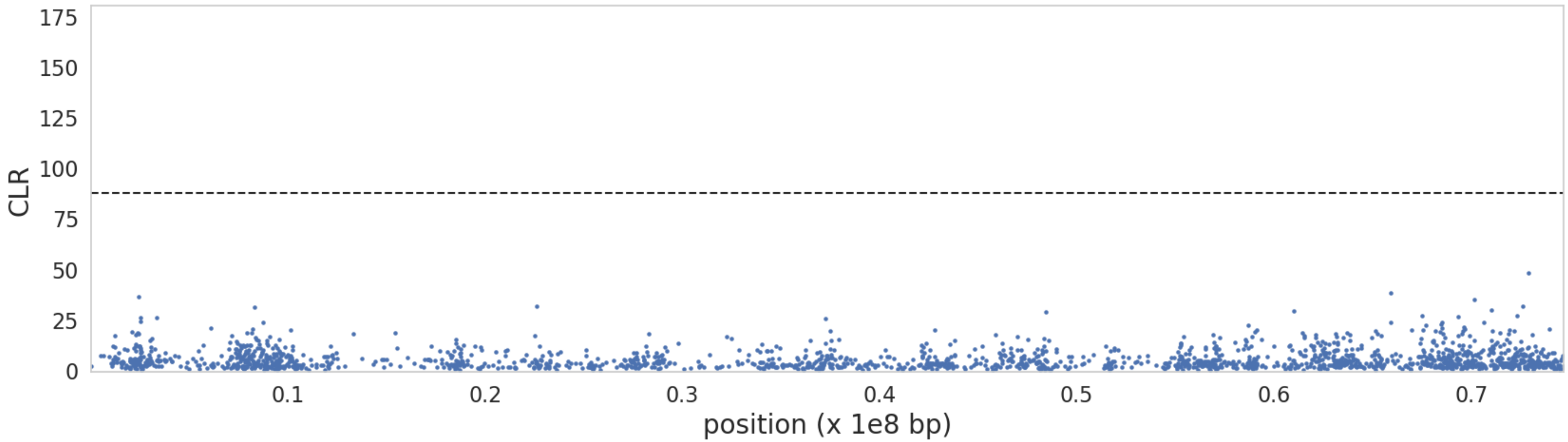


**Supplementary Figure S40:** $B_{0MAF}$ balancing selection scan results for chromosome **17**. The x-axis represents the position along the chromosome, and the y-axis represents the CLR value at each 1kb window (with a 50bp step size). Black dashed line represents the null threshold for balancing selection inference. Candidate genes are labelled on plot.

**Supplementary Figure S41**

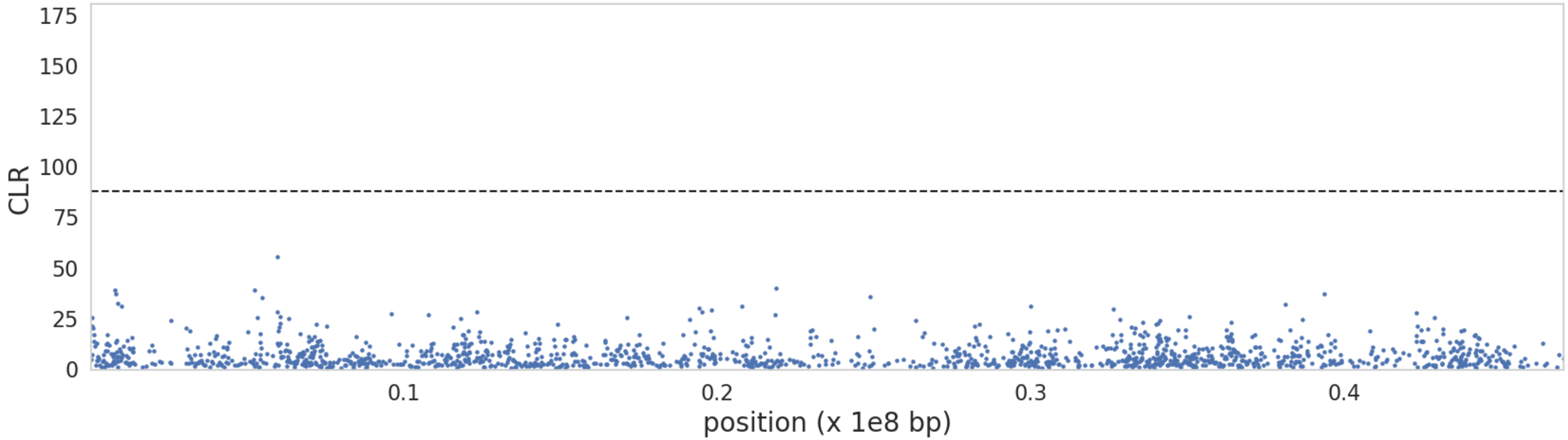


**Supplementary Figure S41:** $B_{0MAF}$ balancing selection scan results for chromosome **18**. The x-axis represents the position along the chromosome, and the y-axis represents the CLR value at each 1kb window (with a 50bp step size). Black dashed line represents the null threshold for balancing selection inference. Candidate genes are labelled on plot.

**Supplementary Figure S42**

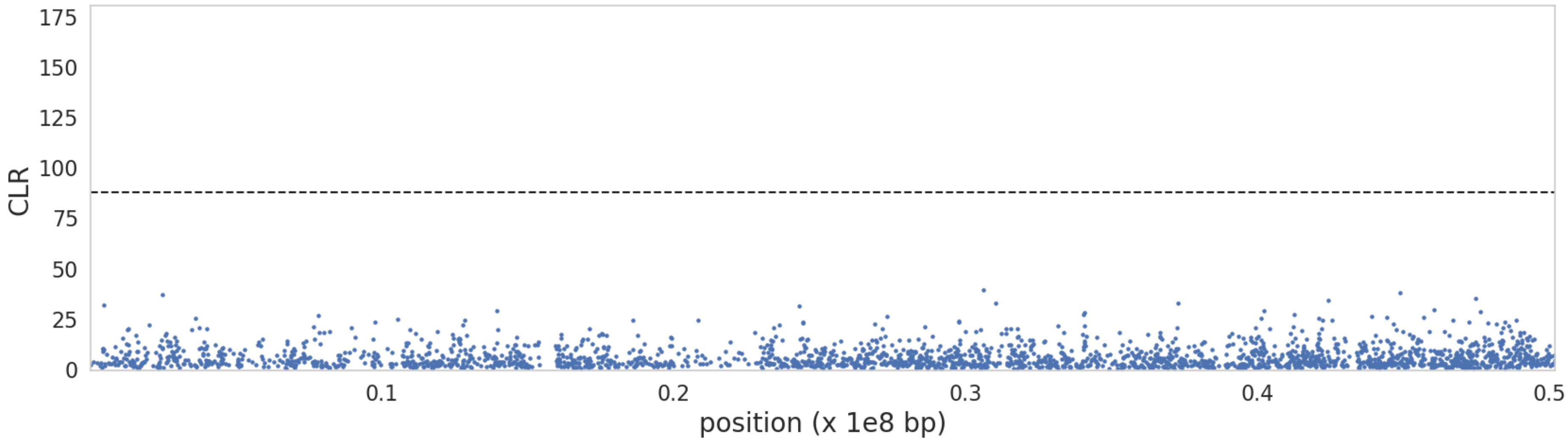


**Supplementary Figure S42:** $B_{0MAF}$ balancing selection scan results for chromosome **19**. The x-axis represents the position along the chromosome, and the y-axis represents the CLR value at each 1kb window (with a 50bp step size). Black dashed line represents the null threshold for balancing selection inference. Candidate genes are labelled on plot.

**Supplementary Figure S43**

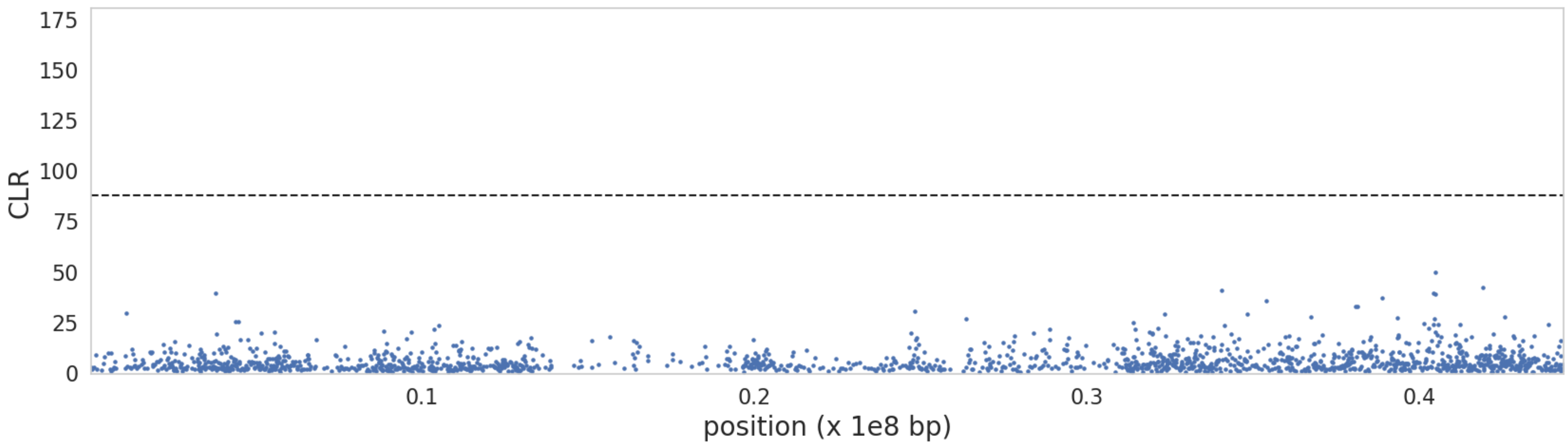


**Supplementary Figure S43:** $B_{0MAF}$ balancing selection scan results for chromosome **20**. The x-axis represents the position along the chromosome, and the y-axis represents the CLR value at each 1kb window (with a 50bp step size). Black dashed line represents the null threshold for balancing selection inference. Candidate genes are labelled on plot.

**Supplementary Figure S44**

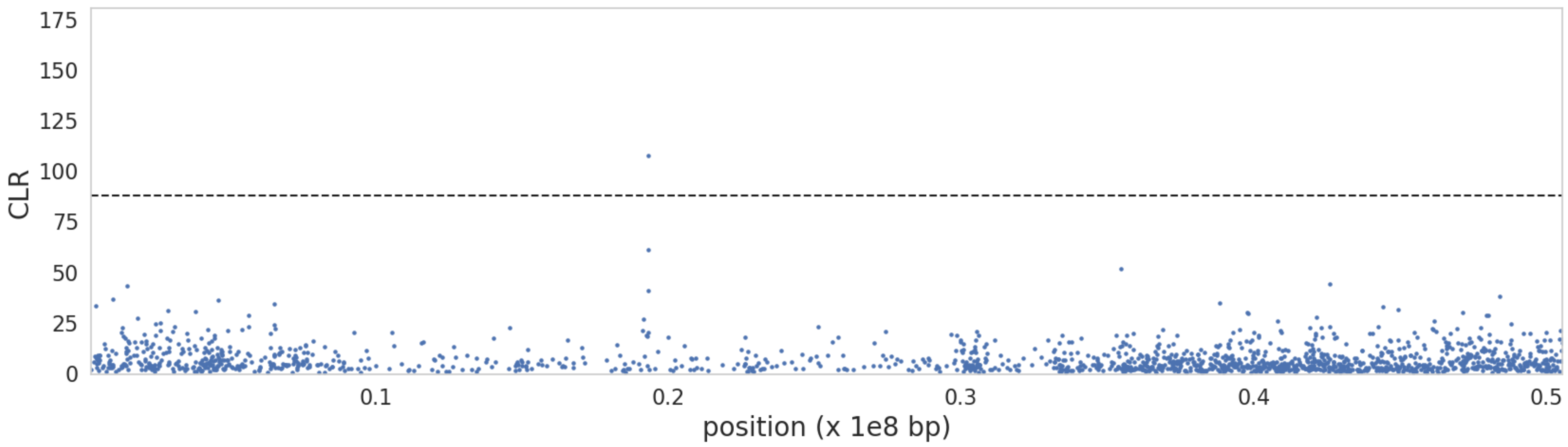


**Supplementary Figure S44:** $B_{0MAF}$ balancing selection scan results for chromosome **21**. The x-axis represents the position along the chromosome, and the y-axis represents the CLR value at each 1kb window (with a 50bp step size). Black dashed line represents the null threshold for balancing selection inference. Candidate genes are labelled on plot.

**Supplementary Figure S45**

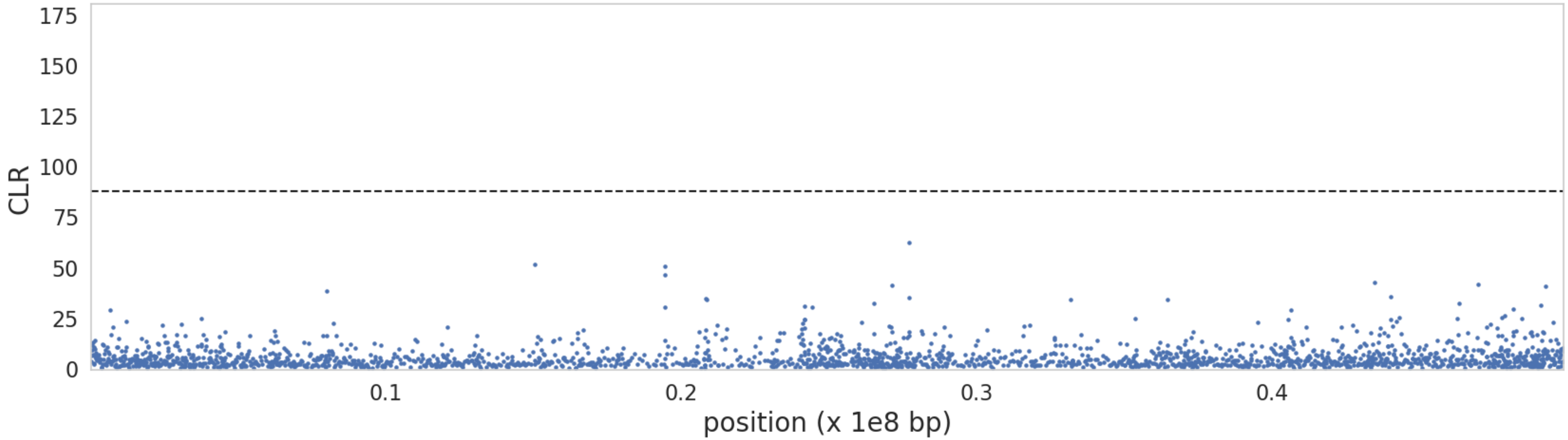


**Supplementary figure S45:** $B_{0MAF}$ balancing selection scan results for chromosome **22**. The x-axis represents the position along the chromosome, and the y-axis represents the CLR value at each 1kb window (with a 50bp step size). Black dashed line represents the null threshold for balancing selection inference. Candidate genes are labelled on plot.
